# Mosquito community composition shapes virus prevalence patterns along anthropogenic disturbance gradients

**DOI:** 10.1101/2021.02.04.429754

**Authors:** Kyra Hermanns, Marco Marklewitz, Florian Zirkel, Anne Kopp, Stephanie Kramer-Schadt, Sandra Junglen

## Abstract

Previously unknown pathogens often emerge from primary ecosystems, but there is little knowledge on the mechanism behind. Most studies analyzing the influence of land-use change on pathogen emergence focus on a single host-pathogen system and often observe contrary effects. We studied virus diversity and prevalence patterns in natural and disturbed ecosystems using a multi-host and multi-taxa approach.

We detected 331 viral sequences pertaining to 49 viruses of ten RNA-virus families. Highest host and virus diversity was observed in pristine and intermediately disturbed habitats. The majority of the viruses was detected with low prevalence. However, nine viruses were found frequently of which five viruses increased in prevalence from pristine to disturbed habitats, in congruence with the dilution effect hypothesis. Interestingly, the observed increased prevalence of these five viruses in disturbed habitats was not caused by higher host infection rates but by increased host abundance, an effect tentatively named abundance effect.

**Impact statement:** Our data show that ecosystem disturbance can lead to a turnover in host community composition and that more individuals of a single species is a key driver of virus emergence.

## Introduction

A major challenge in understanding the emergence of infectious disease is to identify the driving factors responsible for changes in infectious disease dynamics. New infectious diseases mostly emerge in tropical regions which have undergone strong ecological and economic change (1). Tropical rainforests are terrestrial ecosystems with a high biodiversity, constituting a habitat with a diverse range of hosts, vectors and pathogens. This high host richness likely corresponds to a high pathogen richness, as each host is likely to carry its own specific pathogens (2). Pristine rainforests are subject to large scale anthropogenic land use transformation leading to increased contact between humans, wildlife, and pathogens endemic in the wildlife population (3, 4). Disturbed habitats often show a drastic decline in biodiversity or turnover of host species community composition, which is accompanied by an increase of species that are resilient to disturbance, so called generalist species. How these changes in community composition and pathogen abundance influence infection risks of humans and livestock are still unclear and a matter of scientific debate (5).

In this context, the dilution effect hypothesis postulates that a high diversity of different host species dilutes the prevalence of a specific pathogen in intact ecosystems, as the density of competent hosts (hosts that contribute to pathogen transmission and maintenance) is diluted by the presence of non-competent hosts. Biodiversity loss increases the infection prevalence of this specific pathogen in the disturbance-resilient, competent host species (6). Prerequisites for the occurrence of a dilution effect are the presence of species which differ in their susceptibility for a particular pathogen and a lower risk of extinction of competent hosts under habitat disturbance. This may then lead to a higher relative frequency of competent host species in modified habitats (7-9). Hence, in the case of zoonotic pathogens, the disease risk for humans is increased (6, 10). Studies confirming the dilution effect hypothesis focused on pathogens known to cause outbreaks in humans and livestock (e. g. West Nile virus (WNV), Sin Nombre virus and *Borrelia* spp.) and use generalist species as host (11-14). However, it seems that this effect cannot be generalized and that diverse mechanisms regulate infectious disease transmission dynamics in a host-, pathogen- and situation-dependent manner (15, 16). Some studies observed a contrary effect for WNV, Usutu virus and *Borellia* spp., where the infection rate or density of infected hosts increased with biodiversity, referred to as amplification effect (17-19).

These conflicting results indicate that biodiversity loss has complex impacts on disease risk and effects can be heterogeneous and scale-dependent (16, 20-22). Partial views on single host species or single pathogens cannot reveal general mechanisms. Existing studies either focused only on pathogen discovery aside from an ecological context or on one specific pathogen in a specific host across different habitats (12, 13, 17, 23). Studies with a wider scope analyzing community composition of entire host groups and assessing the genetic diversity of their pathogens in undisturbed and disturbed habitats are lacking and would be paramount for a comprehensive understanding of biodiversity-infection relationships. To this end, analyzing species interactions is crucial for a mechanistic understanding of the factors governing emerging infectious diseases (24).

Here, we provide a comprehensive analysis combining fields of community ecology and virology to understand how viral richness depends on host richness, and which role host-habitat associations play in viral prevalence patterns. Several arthropod-borne viruses (arboviruses) are prominent examples for emerging viruses that originated in biodiverse habitats and started from there their spread to new territories where they caused outbreaks and epidemics, e.g. dengue virus (DENV), yellow fever virus (YFV) and Zika virus (ZIKV) (25, 26). Nearly all arboviruses are RNA viruses, which can rapidly evolve facilitating the acquisition of new hosts and geographic spread (27). Mosquitoes can transmit arboviruses during blood-feeding to vertebrates and thereby act as virus vectors. In addition, they can also be infected with their own specific viruses, which cannot be transmitted to vertebrates and are called insect-specific viruses (28). These may serve as models to study effects of land use change and changes in host community composition on mosquito-borne viruses.

In a preliminary analysis we had demonstrated that the prevalence of three insect-specific viruses isolated from mosquitoes sampled along an anthropogenic disturbance gradient in Côte d’Ivoire, West Africa, increased from pristine to disturbed habitat types (29). Subsequently, a plenitude of previously unknown RNA viruses was identified in these samples establishing novel species, genera and even families (30-41). To this end, we established broad-range generic PCR-assays for all viruses identified previously in these samples, as well as for all major virus taxa containing arthropod-associated viruses. With this, we aimed to assess the viral genetic diversity along with the prevalence patterns of each detected virus in an entire family of hosts (*Culicidae*) sampled across habitat types. We expect a higher species diversity in mosquitoes of the family *Culicidae* in habitats of low or no disturbance and predict a turnover in mosquito species composition along the disturbance gradient concomitant with an increase in disturbance-resilient mosquito species and their viruses. We thus hypothesize that habitat perturbation and subsequent changes in community composition influence virus transmission dynamics. This multi-host and multi-taxa study provides insight into common and distinct micro-evolutionary patterns of virus emergence and geographic spread at the interface of pristine and modified landscapes.

## Materials and Methods

### Mosquito collection

In total, 4562 female mosquitoes were collected in five habitat types along an anthropogenic disturbance gradient in the Taï National Park in Côte d’Ivoire in 2004 (29). Habitat types represented pristine forest (PF), secondary forests (SF), agricultural areas (A), camps within primary forest (C) and villages (V) in the order of low to high human disturbance. Mosquitoes were identified morphologically (29) and based on their COI sequences (42). Mosquito heads were homogenized in 430 pools consisting of 1 to 50 individuals according to species and sampling location (29) resulting in 98 pools (764 mosquitoes) from the primary forest, 98 pools (1083 mosquitoes) from the secondary forest, 100 pools (1153 mosquitoes) from agricultural areas, 66 pools (568 mosquitoes) from research camps as well as 68 pools (994 mosquitoes) from two villages.

### RT-PCR screenings and sequencing

RNA was extracted from the pooled supernatants using the QIAamp Viral RNA Mini Kit (Qiagen). The SuperScript™ III Reverse Transcriptase (Invitrogen - Thermo Fisher Scientific, Waltham, USA) was used for cDNA synthesis according to the manufacturer’s instructions. Generic RT-PCR assays were established based on alignments of the RNA-dependent RNA polymerase (RdRp) sequences for the following taxa, peribunyaviruses, jonviruses, feraviruses, rhabdoviruses, flaviviruses, iflaviruses, orbiviruses and Cimodo virus. The RT-PCRs were designed based on sequence information from viruses, which have been previously isolated from these mosquitoes in cell culture (31-40), and from all major arbovirus taxa. Primer sequences and cycling conditions are available upon request. In addition, we used previously described assays for mesoniviruses (40), phenuiviruses (43), flaviviruses (44, 45) and alphaviruses (30) to test the samples. This entire approach allowed the detection of viruses regardless of isolation success in cell culture. PCR products were sequenced by Sanger sequencing (Microsynth AG, Balgach, Switzerland). In addition to the generic assays, specific qPCRs were used to test for Cavally virus, Nsé virus and Mikado virus (40, and see below). The entire RdRp motifs of the third conserved region were amplified from all sequences with more than 5% divergence to another sequence using primer walking. Selected virus isolates were sequenced by metagenomic sequencing as previously described (30).

### Genomic and phylogenetic analyses

All sequences were assembled and analyzed in Geneious R9.1.8 (46). Viral sequences were categorized based on genetic similarity and phylogenetic analysis. Sequences were compared to the NCBI database using blastn and blastx. Sequences with less than 95% amino acid identity to known viruses were considered as putative novel virus species and named Cimo virus (acronym for <u>C</u>ôte d’Ivoire and <u>mo</u>squito) with ascending numbering. Viruses isolated in cell culture and those with completely sequenced genomes received individual names. For phylogenetic analyses, the amino acid sequences (families *Phenuiviridae, Peribunyaviridae, Phasmaviridae, Rhabdoviridae* and *Iflaviridae*) or the nucleotide sequences (genera *Orbivirus, Flavivirus* and *Alphavirus* and family *Mesoniviridae*) of the detected viruses and the related established virus species were aligned by MAFFT-E v7.308 (47) in Geneious. An optimized maximum-likelihood phylogenetic tree with the substitution model based on Smart Model Selection (48) as implemented in PhyML (mainly LG or GTR, respectively) and 1,000 bootstrap replicates was calculated using PhyML (49). For the detected jonviruses the nonsynonymous and synonymous substitution rates were inferred using FEL (Fixed Effects Likelihood) as implemented in Datamonkey (50).

### Virus growth analyses

Viruses were isolated in cell culture as described before (29). For the four novel viruses, Sefomo virus, Mikado virus, Tafomo virus and Sassandra virus, growth kinetics were performed using the mosquito cell line C6/36 (ECACC 89051705, *Aedes albopictus*). C6/36 cells were cultivated and seeded as previously described (51). The cells were infected at a MOI of 0.1 and incubated for 3 days at 28, 30, 32 and 34°C. Every 24 hours 75 µl supernatant was taken and RNA was extracted using the NucleoSpin RNA Virus kit (Macherey-Nagel). cDNA was synthesized using the SuperScript IV reverse transcriptase (Invitrogen – Thermo Fisher Scientific) according to the manufacturer’s instructions. Specific quantitative RT-PCRs were established for all four viruses. Respective primer and probe sequences are SefomoV-F: 5’-TGGTTGAGACCTTCTGAGACTTTTC-3’, SefomoV-R: 5’-CAAAGGCCATCCCGAAGTATC-3’, SefomoV-TM: 5’-6-FAM/CCATTACAC/ZEN/CTCATCCCTATTTCATGCTGG/Iowa Black^®^-FQ-3’, MikadoV-F: 5’-GA-GAACTGTCAAAAATGGAGAAGAGA-3’, MikadoV-R: 5’-AGATGGCACCATTTTCAGTGATATAG-3’, MikadoV-TM: 5’-6-FAM/GCCAACAGC/ZEN/CAATTAAGAGAATGA/Iowa Black^®^-FQ-3’, TafomoV-F: 5’-AATCTGATCTGGAGGACGAGTTG-3’, TafomoV-R: 5’-GCTGTTGATTAGCTGTG-CATGAT-3’, TafomoV-TM: 5’-6-FAM/GGTTCTTGC/ZEN/TGGACCAGGTG/Iowa Black^®^-FQ-3’, SassandraV-F: 5’-CATTTTGGAAGGAGATTTTTCGA-3’, SassandraV-R: 5’-GATCAAATTTCCAA-TAGCCCATAAA-3’ and SassandraV-TM: 5’-6-FAM/TGGACCTCA/ZEN/AGCGGATTCAAC-CGT/Iowa Black^®^-FQ-3’.

### Statistical analysis

#### Virus prevalence

To estimate virus prevalence, the minimum infection rate (MIR) and the maximum likelihood estimation (MLE) per 1000 mosquitoes was calculated with the Excel Add-In PooledInfRate, version 4.0 (52). Virus data analyses were performed in GraphPad Prism 7.04 (GraphPad Software, San Diego, USA).

#### Biodiversity analyses

All biodiversity analyses were done with R version 3.6.2 (53). We used rarefaction curves and Hill numbers of diversity order q = 0 (species richness) to compare mosquito communities between the five habitats ((54); R-package ‘iNEXT’: (55, 56)). To this end, we summed the number of mosquito species per habitat and excluded non-defined species from the analysis. For detection of underlying gradients, we used the Bray-Curtis dissimilarity index ((57); R-package ‘vegan’ (58)) and hierarchical cluster analysis to group main mosquito species according to their habitat association (R-package ‘pheatmap’). To assess whether there are significant associations between these mosquito groups and habitat, we fitted a log-linear model (generalized linear model with ‘log’ link and Poisson error distribution) to the number of mosquitoes counted per group as response variable, with mosquito group and habitat as covariates in interaction. Since there were no counts of *Coquillettidia* species in camp and village, we added a small constant of 1 to all responses to avoid singularities in the model outcome and to be able to model it with a log transformation. We assessed significance of the single terms using a log likelihood ratio test (LRT) of this saturated model.

We used Spearman’s rho on the full dataset to assess which viruses were associated with each other in the different pools. We focused our analyses on abundant viruses we had found at least 10 times in all pools. For assessing virus-habitat relationships expressed as the probability of detecting one of the abundant viruses in a specific habitat, we fitted a generalized linear mixed model with binomial error structure and logit link and the respective virus type detected (1) or not (0) as response. As covariates, we included the number of mosquitoes per pool and the habitat. We included the number of mosquitoes per pool as controlling variable, because the probability to detect any virus in a pool logically increases when more potential hosts are combined into a pool. Pairwise differences in virus infection probabilities in the habitats were assessed post hoc with Tukey contrasts (R-package multcomp).

### Nucleotide Sequence Accession Numbers

The sequence fragments and viral genomes were assigned the GenBank accession numbers … *(pending)*, respectively. Accession numbers of representative sequences are listed in Table 1.

**Table 1:**
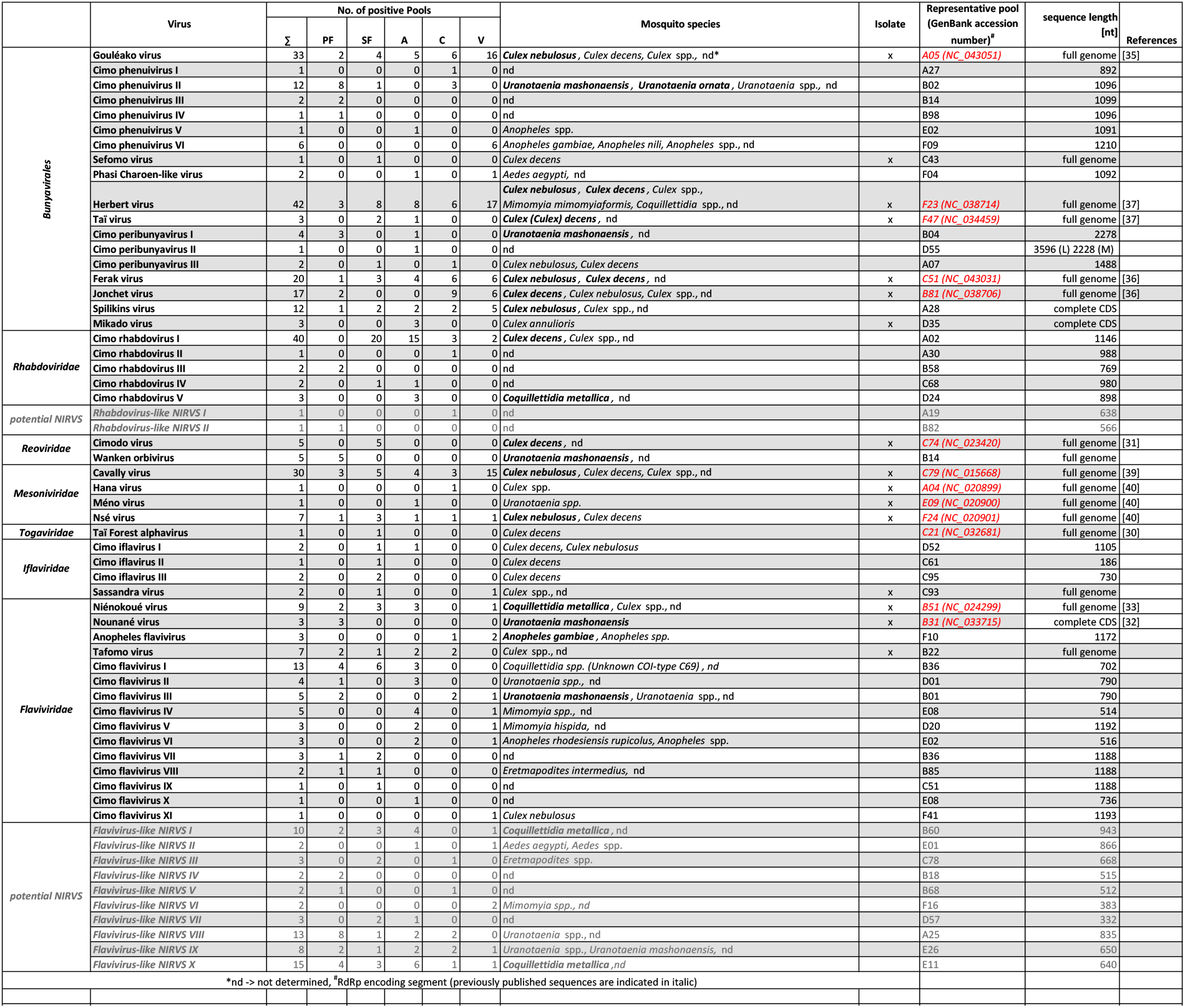
Distribution and host-association of detected viruses. Number of positive pools per habitat and mosquito host species of all detected viruses and virus-like sequences. The main mosquito host species are indicated in bold letters.

## Results

### Biodiversity analyses

The dataset included 42 unique mosquito species units. We counted and rarefied (r given as asymptotic diversity estimate) the highest number of species (n) in the primary forest (PF, n = 20 | r= 38 | with i number of individuals = 462), followed by camp (C, n = 18 | r = 24 | i =418) and secondary forest (SF, n = 14 | r = 24 | i = 651), agriculture (A, n = 17 | r = 18 | i = 882), and village (V, n = 13 | r = 15 | i = 857) (**Fig. 1**).

**Figure 1:**
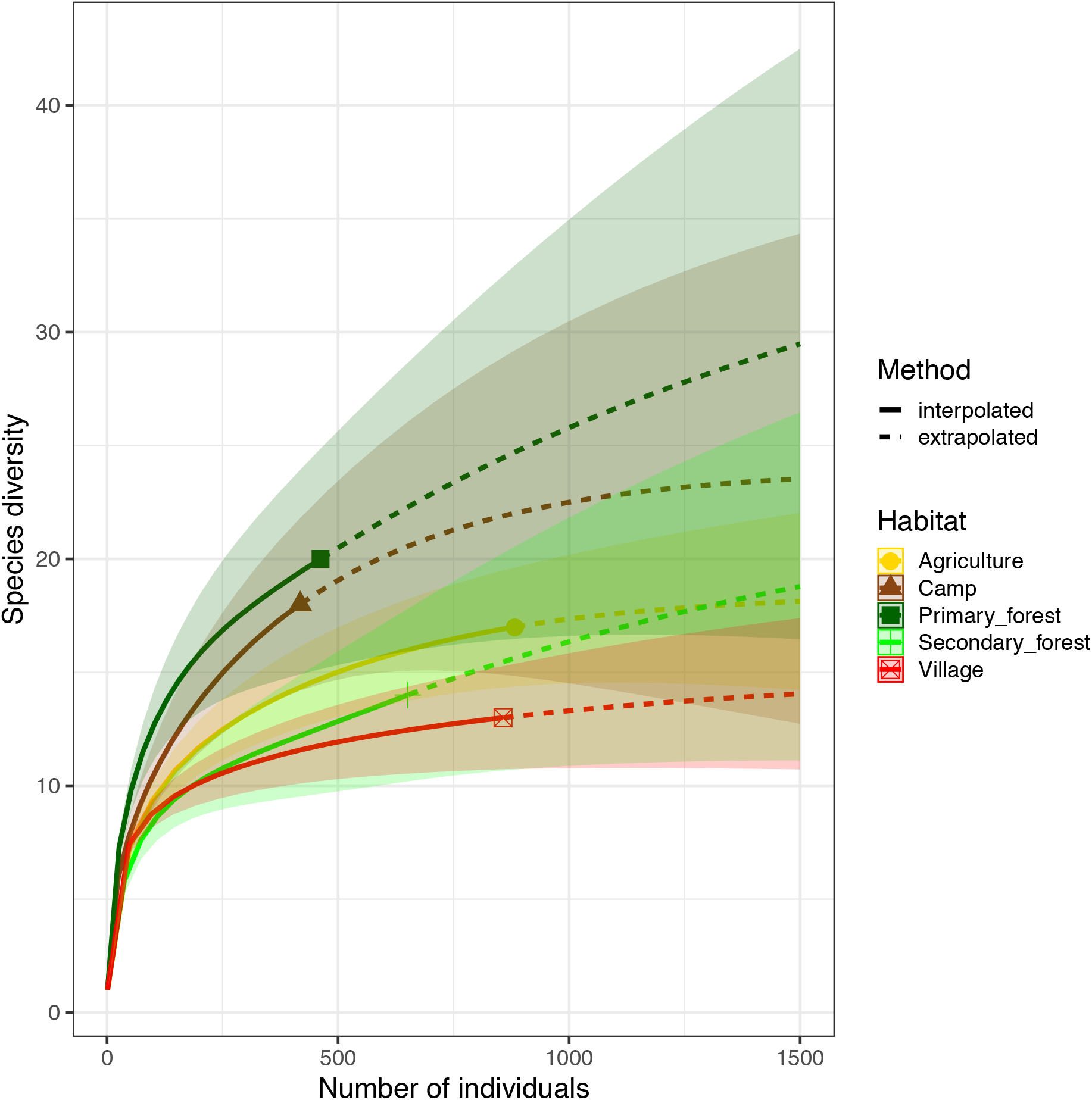
Mosquito species richness per habitat. Rarefaction curves (with 95% confidence intervals) show the observed (solid lines) and interpolated (dotted line) mosquito diversity in the five habitats.

Rarefaction curves were still increasing for camp, primary and secondary forest, indicating that a higher trapping effort would have led to a higher number of species. However, confidence intervals were strongly overlapping, especially for camp and primary forest and for agriculture, secondary forest and village. Bray-Curtis dissimilarity (BCd) yielded the highest difference in community assembly between primary forest and village (BCd = 0.90), and highest similarity between agriculture and secondary forest (BCd = 0.69). The hierarchical cluster analysis confirmed the finding that agriculture and secondary forest clustered together via *Culex decens* and *Culex nebulosus*, while villages and camp mainly clustered via *Culex nebulosus*. Agriculture had high amounts of *Culex annulioris* and *Coquillettidia metallica*, while primary forest mosquito communities were characterized by *Uranotaenia* species. Villages were mainly characterized by *Anopheles* species (**SI Fig. 1 a**). We therefore grouped our mosquitoes to the eight major groups *Culex decens* (i = 1101), *Culex nebulosus* (i = 719), *Culex annulioris* (i = 263), *Culex sp*. (i = 222), *Coquillettidia sp*. (i = 95), *Uranotaenia sp*. (i = 502) and *Anopheles sp*. (i = 294), while all others, mainly the non-defined ones, were termed ‘others’ (i = 1366). Based on these findings, our anthropogenic disturbance gradient was as follows (from low to high disturbance): PF-SF-A-C-V. Significant associations were found between main mosquito groups and habitat (LRT *p* < 0.001, df = 28, deviance = 3782; **SI Table 1 and SI Fig. 1 b**).

### Assessment of the genetic virus diversity

In total, we found 331 viral RdRp sequences pertaining to 34 putative novel viruses and to 15 previously identified viruses of the families *Phenuiviridae, Peribunyaviridae* and *Phasmaviridae* of the order *Bunyavirales*, as well as of the families *Flaviviridae, Togaviridae, Rhabdoviridae, Reoviridae, Iflaviridae* and *Mesoniviridae* (**Table 1**). Sequences with at least 5% pairwise amino acid distance to known viral RdRp sequences were suggested to pertain to distinct viral species.

The family *Phenuiviridae* (order *Bunyavirales*) includes important arboviruses within the genus *Phlebovirus* but also numerous insect-specific viruses that for example belong to the genera *Goukovirus* and *Phasivirus* (59). Nine distinct phenuiviruses were detected, which included seven novel viruses, named Sefomo virus (acronym for <u>se</u>condary <u>fo</u>rest <u>mo</u>squito virus) and Cimo phenuivirus I-VI, as well as Gouléako virus (GOLV) and Phasi Charoen-like virus (PCLV) (35, 60). For all viruses, the number of positive samples per habitat is summarized in **Table 1**. Phylogenetic analyses showed that the viruses grouped with insect-specific viruses of the genera *Goukovirus, Phasivirus, Hudivirus* and *Beidivirus*, as well as with the unclassified insect viruses related to the uncultured virus isolate acc 9.4 (**Fig. 2a**). Interestingly, Cimo phenuivirus V branched basal to tenuiviruses that are transmitted between plants by planthoppers (61). Cimo phenuivirus V was found in *Anopheles* spp. mosquitoes collected at a rice plantation. At this point we cannot differentiate whether the mosquito ingested infected plant material or if this novel tenui-like virus can infect mosquitoes.

**Figure 2:**
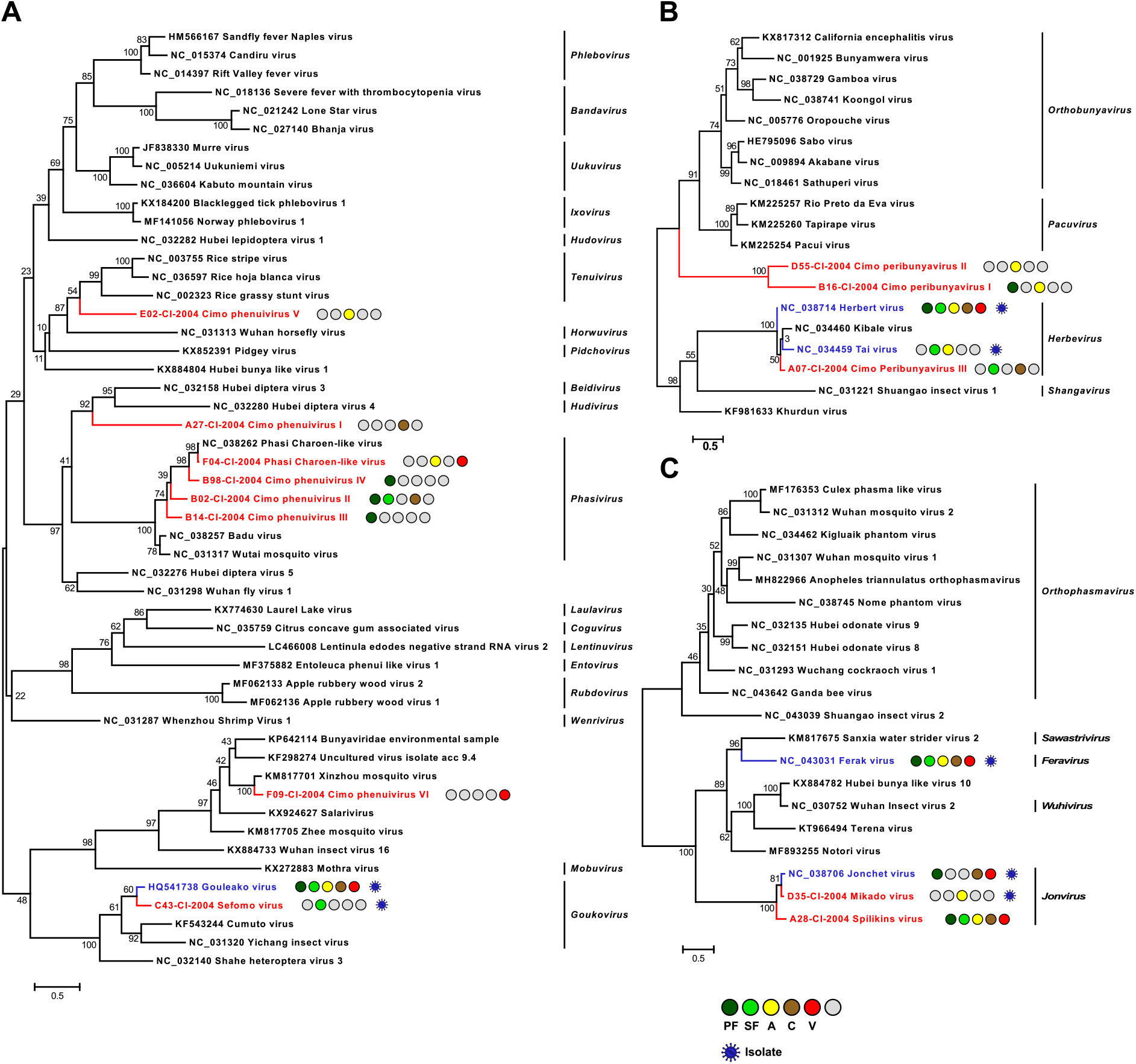
Phylogenetic analyses of detected bunyaviruses. Phylogenetic trees were inferred with PhyML (LG substitution model) based on MAFFT-E protein alignments covering the conserved RdRp motifs of the families *Phenuiviridae* (**A**), *Peribunyaviridae* (**B**) and *Phasmaviridae* (**C**). Novel viruses from this study are indicated in red and previously published viruses detected in our data set are indicated in blue. Sample origin from the different habitat types is indicated by colored circles while no detection is indicated by grey circles. Live virus isolates are marked with a blue virion. Abbreviations are PF, primary forest; SF, secondary forest; A, agriculture; C, camp and V, village.

The family *Peribunyaviridae* (order *Bunyavirales*) consists of two arbovirus genera (*Orthobunyavirus* and *Pacuvirus*) and two genera which contain insect-restricted viruses (*Herbevirus* and *Shangavirus*) (62). We identified two novel peribunyaviruses, named Cimo peribunyavirus I and II. These viruses formed a monophyletic clade that shared a most recent common ancestor with arboviruses of the genera *Orthobunyavirus* and *Pacuvirus* (**Fig. 2b**). In addition, the insect-specific herbeviruses Taï virus and Herbert virus (HEBV) (37), as well as a previously undescribed herbevirus, named Cimo peribunyavirus III, were detected that fell into the clade of herbeviruses (**Fig. 2b**).

The family *Phasmaviridae* (order *Bunyavirales*) comprises only insect-specific viruses (59). We found two prototype species of the genera *Feravirus* and *Jonvirus*, Ferak virus (FERV) and Jonchet virus (JONV), which were previously isolated in cell culture from these mosquitoes (36). While a great diversity of novel phasmaviruses has been found since the first discovery of this family, no additional members of the genus *Jonvirus* have been identified (59). Here, we further detected two previously unknown jonviruses, named Mikado virus and Spilikins virus, which are closely related to the prototype virus JONV in phylogenetic analyses (**Fig. 2c**). For both novel jonviruses, the complete coding sequence (CDS) was sequenced and phylogenies of the M and S segments are shown in the **Supplementary Information (SI) Figure 2**. In all phylogenies, the two novel viruses cluster well supported with JONV.

The family *Rhabdoviridae* (order *Mononegavirales*) is highly diversified and currently contains 20 genera. Rhabdoviruses infect vertebrates, arthropods and plants (63). We detected five previously unknown rhabdoviruses, named Cimo rhabdovirus I-V that clustered with different unclassified clades of mosquito-associated rhabdoviruses across the rhabdovirus phylogeny (**Fig. 3a**).

**Figure 3:**
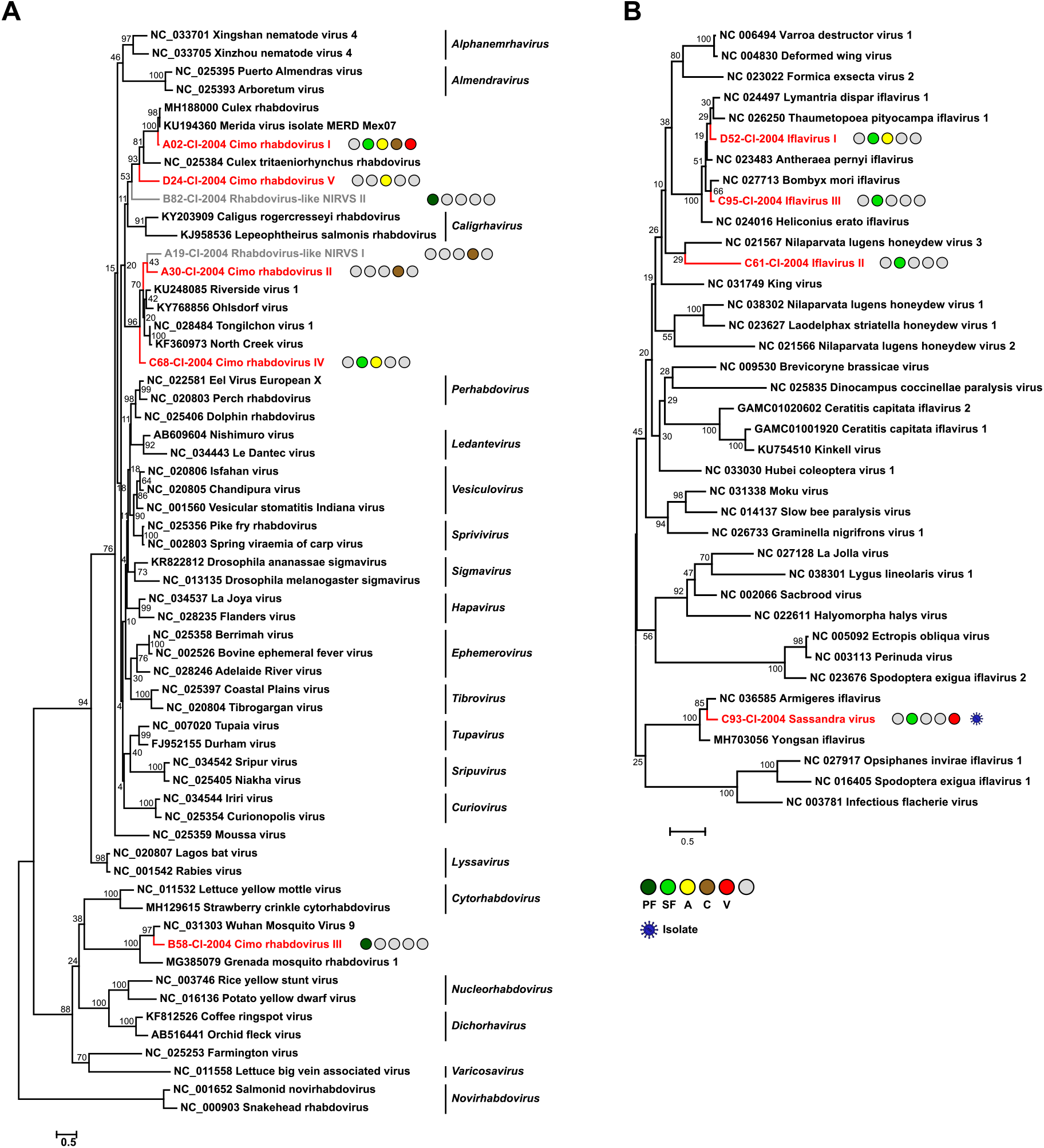
Phylogenetic analyses of detected rhabdoviruses and iflaviruses. Phylogenetic trees were inferred with PhyML (LG substitution model) based on MAFFT-E protein alignments covering the conserved RdRp motifs of the families *Rhabdoviridae* (**A**) and *Iflaviridae* (**B**). Novel viruses from this study are indicated in red and detected virus-like sequences are indicated in grey. Sample origin from the different habitat types is indicated by colored circles while no detection is indicated by grey circles. Live virus isolates are marked with a blue virion. Abbreviations are PF, primary forest; SF, secondary forest; A, agriculture; C, camp and V, village.

The family *Iflaviridae* (order *Picornavirales*) consists of a single genus whose members are restricted to arthropod hosts (64). We detected four novel iflaviruses, named Sassandra virus and Cimo iflavirus I-III, which were placed in three different clades in phylogenetic analyses (**Fig. 3b**). Cimo iflavirus I and III grouped with Bombyx mori iflavirus and other lepidopteran iflaviruses, while the short sequence fragment of Cimo iflavirus II did not form a well-supported clade with known iflaviruses. Sassandra virus clustered with two previously described iflaviruses from mosquitoes (**Fig. 3b**).

The genus *Flavivirus* (family *Flaviviridae*) includes important arboviruses as well as viruses with a single host tropism for arthropods or vertebrates (65). Insect-specific flaviviruses can be divided into two groups. Classical insect-specific flaviviruses form a monophyletic clade in basal phylogenetic relationship to all other flaviviruses while dual-host affiliated insect-specific flaviviruses are phylogenetically affiliated with the arboviruses of this genus (28). We found twelve undescribed flaviviruses, named Tafomo virus (acronym for <u>Ta</u>ï <u>fo</u>rest <u>mo</u>squito virus) and Cimo flavivirus I-XI, as well as a strain of Anopheles flavivirus (66). All sequences clustered within the clade comprising the classical insect-specific flaviviruses in phylogenetic analysis (**Fig. 4a**). In addition, two previously characterized flaviviruses, the dual-host affiliated insect-specific flavivirus Nounané virus and the classical insect-specific flavivirus Niénokoué virus (NIEV), were detected in the mosquitoes (32, 33).

**Figure 4:**
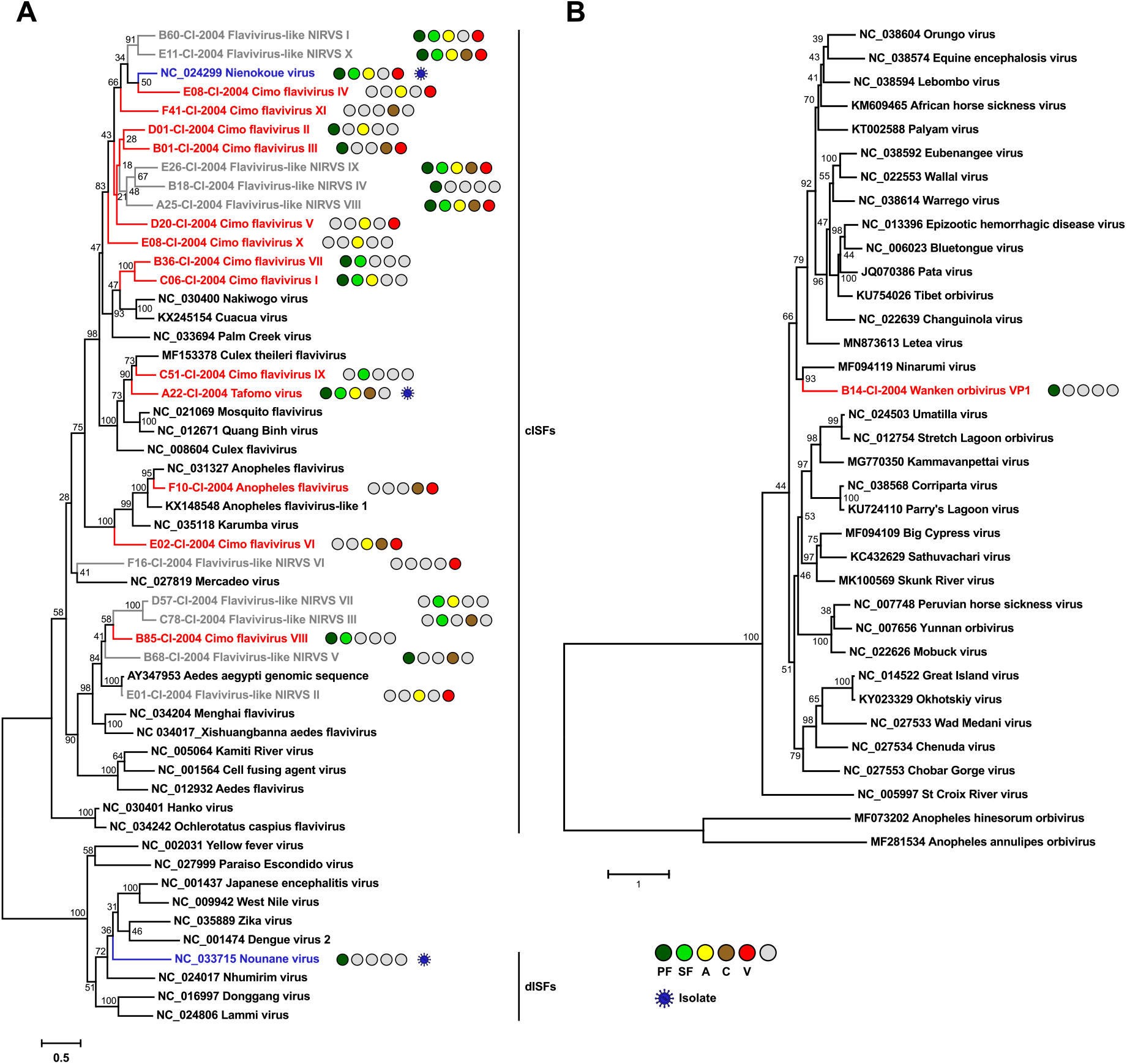
Phylogenetic analyses of detected flaviviruses and orbiviruses. Phylogenetic trees were inferred with PhyML (GTR substitution model) based on MAFFT-E nucleotide alignments covering the conserved RdRp motifs of the genera *Flavivirus* (**A**) and *Orbivirus* (**B**). Novel viruses from this study are indicated in red, previously published viruses detected in our data set are indicated in blue and detected virus-like sequences are indicated in grey. Sample origin from the different habitat types is indicated by coloured circles while no detection is indicated by grey circles. Live virus isolates are marked with a blue virion. Abbreviations are PF, primary forest; SF, secondary forest; A, agriculture; C, camp and V, village.

The family *Reoviridae* comprises 15 genera with highly variable biological properties. Three genera contain arboviruses (*Orbivirus, Coltivirus* and *Seadornavirus*) while viruses belonging to other genera infect vertebrates, plants, fungi or insects (67). One novel orbivirus, named Wanken orbivirus (WKOV), was detected in the mosquitoes. The complete genome of WKOV was sequenced and the polymerase encoded on the first segment showed 56% pairwise amino acid identity to Ninarumi virus. Phylogenetic analysis of the polymerase (VP1) placed WKOV on a branch with Ninarumi virus, a virus detected in *Aedes fulvus* mosquitoes collected in Peru (68). This clade was placed basal to the clade of *Culicoides*- and *Phlebotominae*-transmitted orbiviruses (e.g. Bluetongue virus) and Letea virus, a virus detected in snakes (*Natrix natrix*) captured in Romania (69) (**Fig. 4b**). Additional phylogenetic analyses based on VP3, VP4, VP5 und VP7 also placed WKOV together with Ninarumi virus and Letea virus basal to the clade of *Culicoides*- and *Phlebotominae*-transmitted orbiviruses (**SI Fig. 3**). Additionally, Cimodo virus, which most likely defines a novel reovirus genus and which was previously characterized (31), was found (**SI Fig. 4a**).

Most viruses of the genus *Alphavirus* (family *Togaviridae*) are arboviruses. Contrary to the genus *Flavivirus*, only few insect-specific alphaviruses have been discovered thus far (70). We detected the previously characterized insect-restricted Taï Forest alphavirus (30) but no further alphaviruses were found (**SI Fig. 4b**).

The family *Mesoniviridae* (order *Nidovirales*) consists of insect-specific viruses (40, 71). The four previously described mesoniviruses Cavally virus (CAVV), Nsé virus, Hana virus, and Méno virus were detected in the mosquitoes (40) (**SI Fig. 4c**).

Besides the 12 previously published virus isolates from this sampling (31-41), Sefomo virus, Mikado virus, Sassandra virus, and Tafomo virus were isolated in C6/36 mosquito cells indicating the detection of functional viruses. The viruses replicated well in C6/36 cells incubated at 28-30°C but virus growth was impaired at 32°C for all but Mikado virus. No virus could replicate at 34°C indicating the detection of insect-restricted viruses (**Fig. 5**).

**Figure 5:**
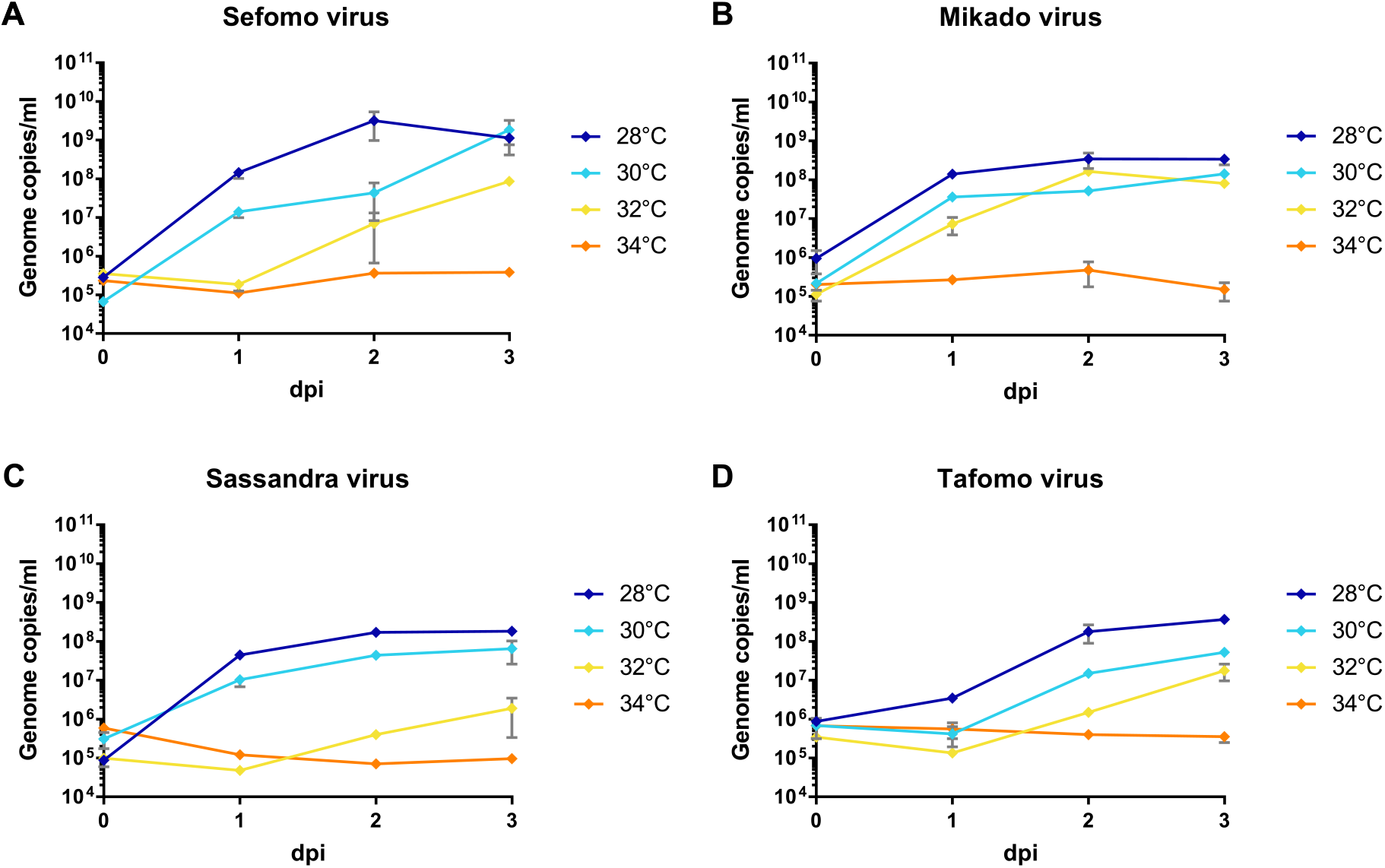
Temperature-dependent replication of novel virus isolates. C6/36 cells were infected with an MOI of 0.1 with Sefomo virus (**A**), Mikado virus (**B**), Sassandra virus (**C**) and Tafomo virus (**D**) in duplicates. Replication was measured for 3 dpi at 28, 30, 32 and 34°C.

### Detection of non-retroviral integrated RNA virus sequences

Genome fragments of flavi-, bunya- and rhabdoviruses were found to have integrated into mosquito genomes and persist as so called non-retroviral integrated RNA virus sequences (NIRVS) (44, 72-74). We detected flavi- and rhabdovirus-like sequences with defective ORFs within the conserved region of the RdRp gene suggesting the detection of NIRVS. The two potential rhabdovirus-like NIRVS encoded either an internal stop codon (rhabdovirus-like NIRVS II – B82) or sequence elongation attempts resulted in rhabdovirus-like sequences that contained frame shifts (rhabdovirus-like NIRVS I – A19). We were further able to amplify these defective virus-like sequences directly from nucleic acid extracts without prior cDNA synthesis suggesting that these sequences were present as DNA. No amplificates were obtained for the sequences with contiguous ORFs (**SI Fig. 5a**).

Furthermore, we obtained ten flavivirus-like sequences with frame shifts, internal stop codons, deletions or integrations (**SI Fig. 5b**). In case of flavivirus-like NIRVS II and III, parts of the potentially integrated sequences were related to sequences from insects including genome loci from *Aedes* mosquitoes. Flavivirus-like NIRVS II was amplified from an *Aedes aegypti* pool (E01) and the integrated fragment (68 nt) consisted of 56 nt with 81% identity to *Aedes aegypti* steroid hormone receptor homolog (AaHR3-2) gene and a 12 nt duplication of the sequence immediately adjacent to the integration site. In addition to the interrupted flavivirus-like sequence in pool E01, an identical continuous flavivirus sequence was obtained from the same pool. A similar observation was made with flavivirus-like NIRVS I (pool B60). The sequence of flavivirus-like NIRVS III (pool C78) was detected in *Eretmapodites intermedius* and undetermined mosquitoes. This sequence was profoundly defective (frameshifts, internal stop codon and deletions) and continued into a 416 nt long sequence with low similarity to *Aedes albopictus* and *Apis* spp. genome loci (**SI Fig. 5b**). The very few and short sequences of *Eretmapodites* spp. available in GenBank most likely impeded the identification of the host genome sequence of flavivirus-like NIRV III.

### Detected viruses show a high host specificity

We next sought to identify if the detected viruses were associated with specific mosquito host species. In total, 43% of the pools were found positive for any virus. The gross majority of the viruses (n= 39) was detected in a single mosquito species and only 10 viruses were detected in two or three different closely related mosquito species which belonged to the same mosquito genus indicating a high host specificity (**Table 1**). Unfortunately, ordination techniques like non-metric multidimensional scaling did not converge due to too few positive findings of viruses in the pools (data not shown). We often detected multiple viruses in one mosquito pool, especially in *Culex* pools, indicating possible co-infections mainly between GOLV, HEBV, CAVV and FERV. These viruses were all associated with *Culex nebulosus* as main mosquito host and consequently often found together in pooled mosquitoes of this species. We found significant positive associations between GOLV and HEBV, as well as between GOLV and CAVV (**Fig. 6**). The association between GOLV and HEBV strongly increased (Spearman’s rho = 0.84, p < 0.05) when considering only the host mosquito *Culex nebulosus*. Here, also correlations of both viruses with FERV and Spilikins virus increased (rho ∼ 0.45, p < 0.01; results not shown).

**Figure 6:**
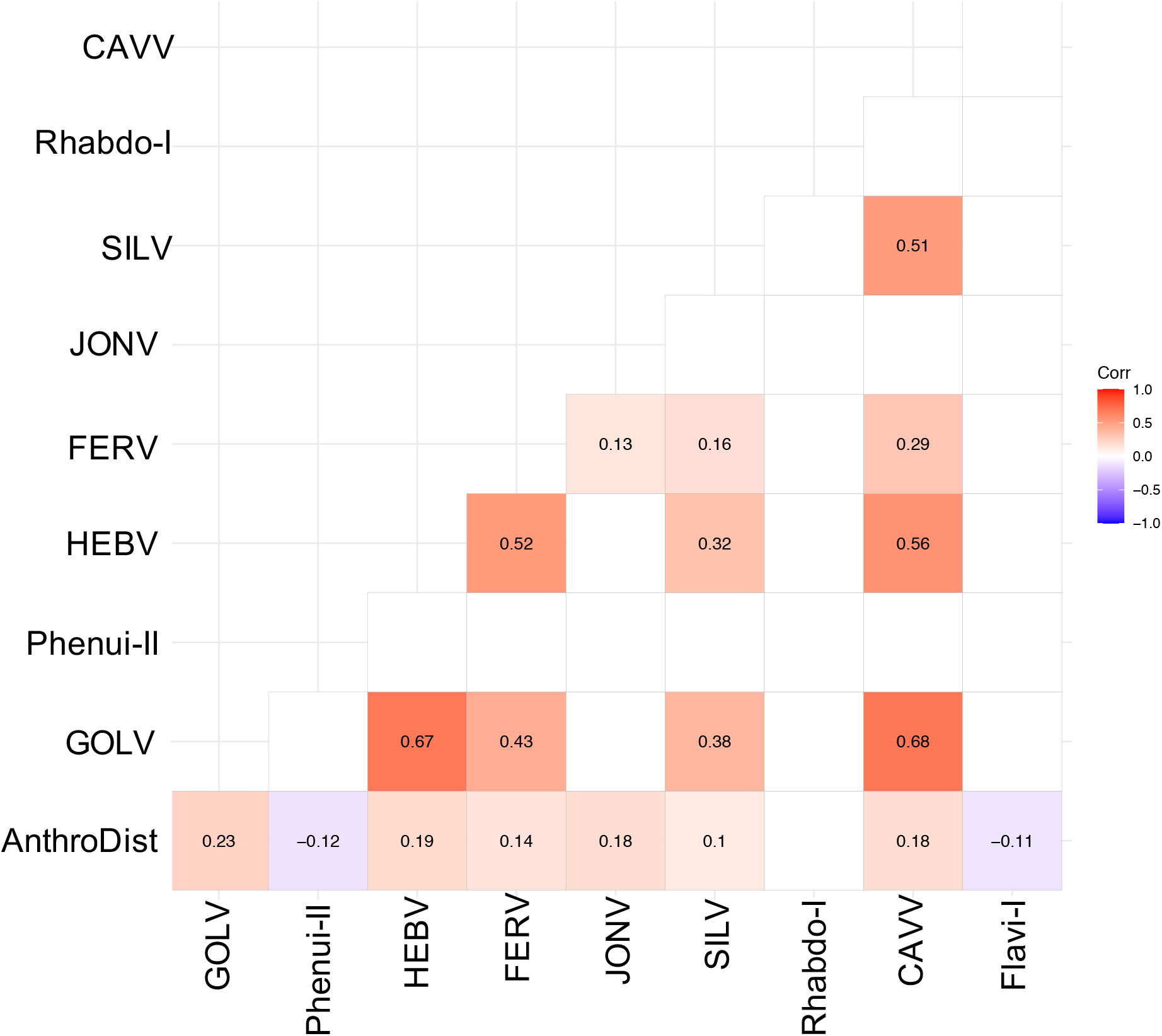
Spearman’s rank correlation rho for the most abundant viruses. Only significant correlations are shown. AnthroDist refers to the gradient of anthropogenic disturbance from low to high: PF, primary forest; SF, secondary forest; A, agriculture; C, camp, and V, village.

Whether these mixed infections result from multiple infected individuals or from co-infected single mosquitoes could not be analyzed, as no homogenates of individual specimens were available. All four viruses could be isolated together in cell culture from multiple pools. The replication of all four viruses in co-infected cell cultures was confirmed by quantitative real-time PCR suggesting no general inhibitory effect and the possibility of simultaneous infection of a single mosquito.

The three jonviruses, JONV, Spilikins virus and Mikado virus, were each also associated with specific mosquito species, namely with *Culex decens, Culex nebulosus* and *Culex annulioris*, respectively. Interestingly, their partial RdRp sequences showed a high degree of variation in the third codon positions which did not alter the translated protein sequences (called synonymous substitutions). For example, JONV and Mikado virus diverged by approximately 20% in their nucleotide (nt) sequences but only by 7% in their amino acid (aa) sequences. Similarly, Spilikins virus and JONV showed nt and aa divergences of 25% and 12%-15%, respectively. FEL analysis of 31 detected jonvirus sequences found significant (p-value <0.05) evidence of negative selection for 88 out of 117 codons. These findings could point towards adaptation to a specific mosquito host under purifying selection. Similar strains with mainly variation in the third codon position were also observed for two Cimodo virus strains, HEBV and Cimo peribunyavirus III, two strains of Cimo peribunyavirus I, two strains of Anopheles flavivirus, as well as Cimo flavivirus I and Cimo flavivirus VII.

### Virus prevalence patterns along the anthropogenic disturbance gradient

According to current hypotheses in infectious disease ecology, ecological perturbation and changes in community composition are expected to influence transmission dynamics of infectious diseases (3, 5, 13). We thus next analyzed the prevalence patterns of all detected viruses as well as abundance of their mosquito host species along the anthropogenic disturbance gradient. The abundance of different mosquito genera varied considerably across the different habitats, as described previously by Junglen et al. (29). The dominant genus across all habitats except in the primary forest was *Culex* (50.5%), whereas in the primary forest mosquitoes of the genus *Uranotaenia* were most frequently sampled. *Culex decens* was most abundant in the secondary forest and agricultural areas, whereas *Culex nebulosus* mosquitoes were the most abundant species in villages and at camp sites located in the primary forest. A large fraction of 56.7%, 43.9% and 52.1% of the sampled mosquitoes in the primary and secondary forest as well as at the camp sites, respectively, could not be identified to species level either due to morphological damage or to limitations of taxonomic keys. The actual number of different species in these habitats is therefore likely to be higher and the relative abundance of some mosquito species may have been underestimated.

The highest virus richness was observed in the intermediately disturbed habitats secondary forest and agriculture (**Fig. 7a**). However, the proportional number of tested mosquitoes was lower in the primary forest so that the richness in this habitat is likely underestimated.

**Figure 7:**
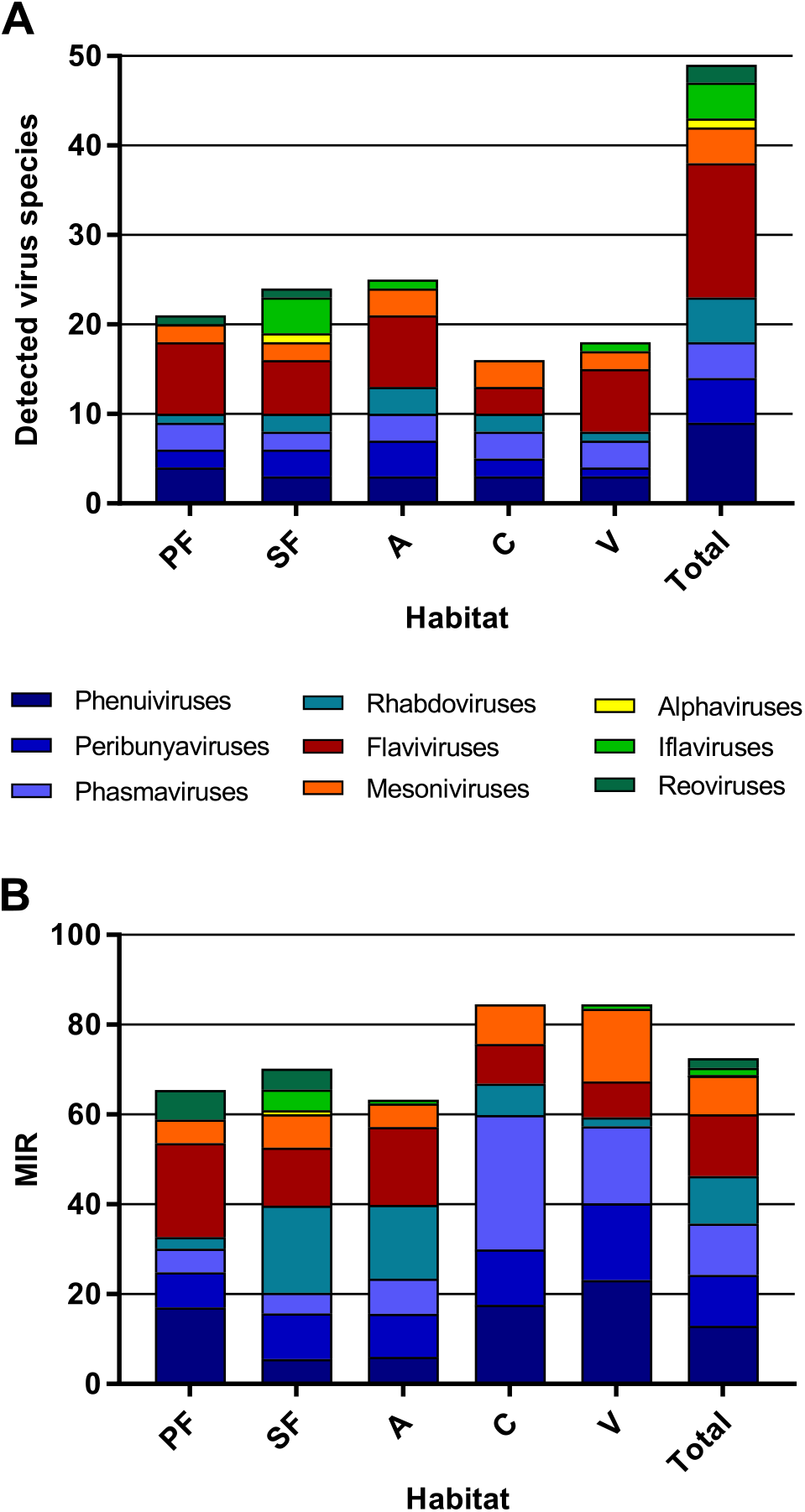
Richness and cumulative MIR across all tested virus taxa. The number of distinct viruses (**A**) and the cumulated MIR per 1000 mosquitoes (**B**) were calculated for all habitat types and for the complete data set. Different virus taxa are shown in different colours.

The majority of the viruses (82%, n = 40) occurred with a low frequency of less than ten positive samples in total (see **Table 1**). The cumulated MIR for all detected viruses was slightly higher in villages and at the camp sites compared to the other habitats (**Fig. 7b**). This effect was mainly caused by the increasing prevalence of several bunyaviruses and one mesonivirus (**SI Fig. 6a, b, f and h**), while other taxa like reoviruses and iflaviruses were found in specific habitats (**SI Fig. 6c and i**), or like flaviviruses and rhabdoviruses increased in prevalence towards pristine or intermediately disturbed habitats (**SI Fig. 6 e and g**). Of note, no amplification effect was observed in habitats with higher biodiversity (**Fig. 7a and b**).

However, nine viruses were found more frequently with detection rates ranging from 12 to 42 strains (**Table 1**). As the sample set consisted of pooled mosquito specimens, we estimated viral infection rates in the different habitats using the two approaches MIR and MLE which showed similar results (**Fig. 8**). Four bunyaviruses (GOLV, HEBV, FERV and Spilikins virus) and one mesonivirus (CAVV), all mainly associated with *Culex nebulosus*, increased in prevalence towards disturbed habitat types (**Fig. 8a, b, c and e - left graphs and 8h**). A trend, that seemed to support the dilution effect hypothesis. In contrast, Cimo rhabdovirus I had its highest prevalence in the intermediately disturbed habitats (secondary forest and agricultural areas) (**Fig. 8d – left graph**) and two viruses increased in prevalence towards the primary and secondary forest (Cimo phenuivirus II and Cimo flavivirus I) (**Fig. 8f and i**). JONV prevalence slightly increased in the villages and more prominently at the camp sites compared to the other habitats (**Fig. 8g**). The prevalence patterns corresponded for eight of the nine viruses to the relative abundance of the main mosquito host species (**Fig. 8**). JONV was the only virus that showed no relationship between mosquito host abundance and virus prevalence suggesting that other factors may influence JONV abundance. Cimo rhabdovirus I and JONV were both mainly associated with *Culex decens* mosquitoes as host. However, in contrast to GOLV, HEBV, CAVV and FERV, which were frequently detected together in *Culex nebulosus* mosquitoes, Cimo rhabdovirus I and JONV were only twice found together in the same pool. This could hint to a possible interference between these viruses and might be a reason for the unusual prevalence pattern of JONV.

**Figure 8:**
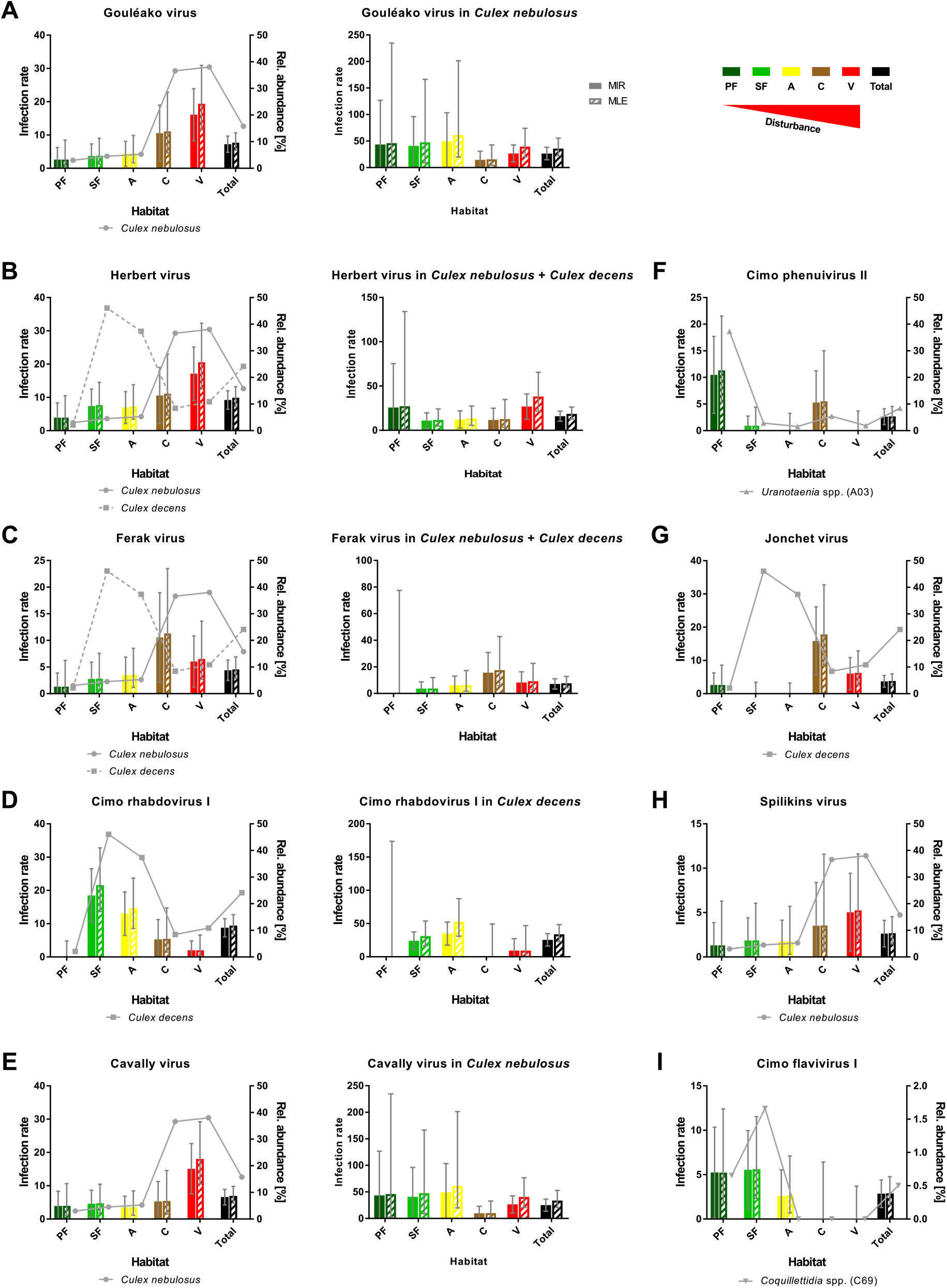
Prevalence patterns of selected viruses along the disturbance gradient. For all viruses, that were detected in >10 pools, GOLV (**A**), HEBV (**B**), FERV (**C**), Cimo rhabdovirus I (**D**), CAVV (**E**), Cimo phenuivirus II (**F**), Jonchet virus (**G**), Spilikins virus (**H**) and Cimo flavivirus I (**I**), the MIR and MLE per 1000 mosquitoes of the whole data set was calculated for all habitat types. The abundance of the main mosquito host species was plotted. The five viruses GOLV (**A**), HEBV (**B**), FERV (**C**), Cimo rhabdovirus I (**D**), and CAVV (**E**) occurred frequently enough in their main mosquito host species (>10 positive pools) to analyze their prevalence in these species. For these viruses, the MIR and MLE per 1000 mosquitoes of the respective species was calculated for all habitat types (right graphs). Significant differences in the infection probability with the most abundant viruses in the different habitats are shown in SI Table 2.

The probability of finding an infection with GOLV, HEBV and CAVV was significantly higher in villages (p < 0.1, **SI Table 2**). These viruses were mainly associated with *Culex nebulosus* mosquitoes. Cimo phenuivirus II was associated with *Uranotaenia* sp. mosquitoes in the primary forest (p < 0.1, **SI Table 2**), whereas JONV and FERV mainly occurred in the camps and villages, associated with *Culex nebulosus, Culex decens* and other mosquito species. Cimo flavivirus I did not show a clear tendency, but mainly occurred in the primary and secondary forest as well as the agricultural sites in *Coquillettidia* sp. and other mosquito species. Cimo rhabdovirus I was almost exclusively associated with *Culex decens* and the detection probability significantly increased in the secondary forest and the agricultural sites (p < 0.1, **SI Table 2**).

We next investigated infection rates only in the main mosquito host species across the different habitat types. Only five of the nine viruses were detected frequently enough in their main mosquito host species (n>10) to calculate host-specific infection rates. Surprisingly, no trend of increasing or decreasing virus prevalence was detected along the disturbance gradient (**Fig. 8a-e – right graphs**) as would be expected in case of a dilution or amplification effect (13, 17). Viral infection rates did not change considerably in their mosquito hosts between disturbed and undisturbed habitat types. Notably, the increase in prevalence of specific viruses resulted from shifts in the mosquito community composition along the gradient, which caused increased abundance rates of the main mosquito host species and concomitantly higher prevalence rates of the viruses they were carrying. We thus refer to this observation as abundance effect (summarized in **Fig. 9**).

**Figure 9:**
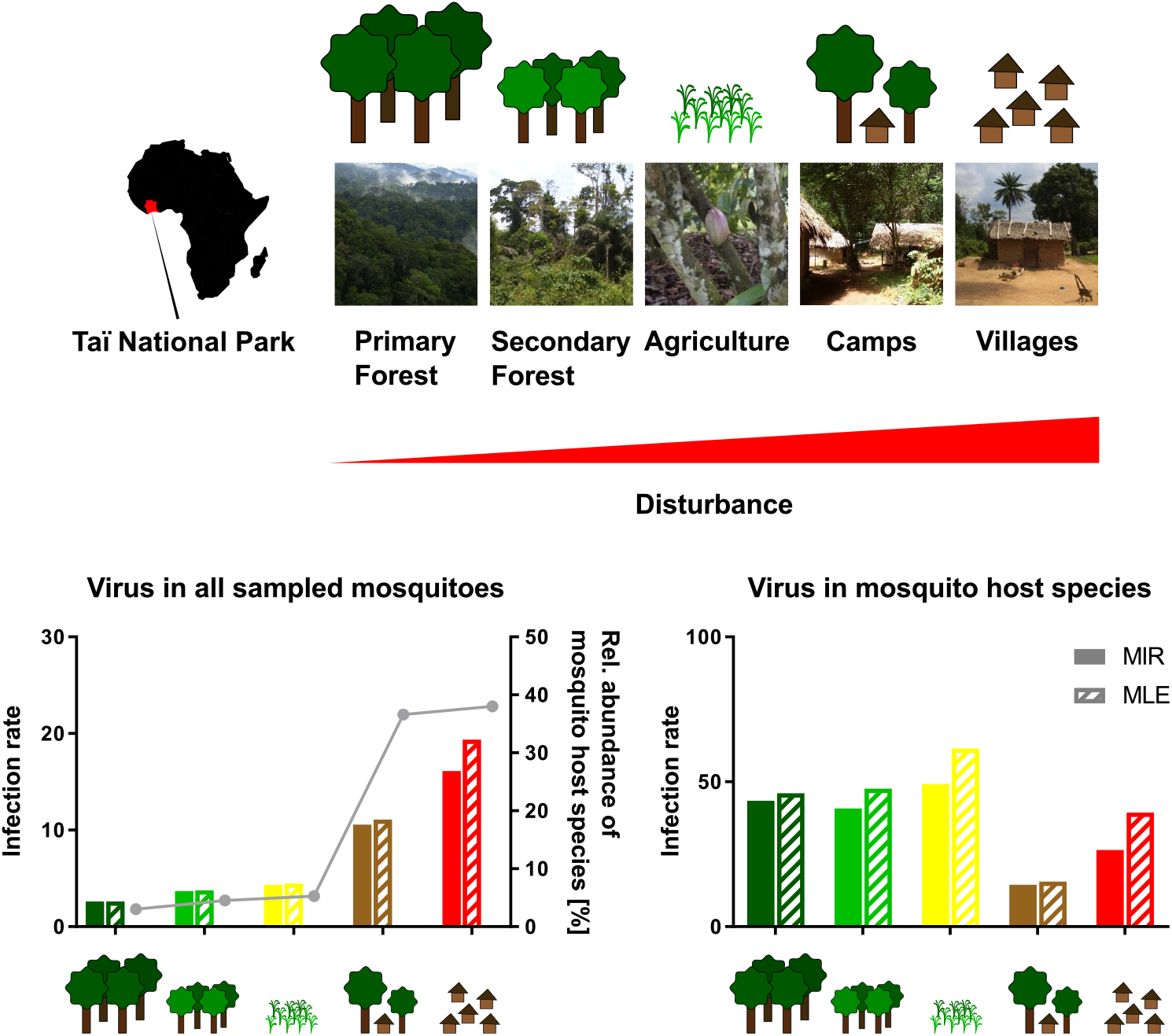
Schematic presentation of the abundance effect. The different habitat types along the anthropogenic disturbance gradient are depicted by photos and drawings. Infection rates are shown for GOLV in all sampled mosquitoes (representing MIR and MLE values as shown in Figure 8) and only in the main mosquito host species, *Culex nebulosus*. The abundance of the main mosquito host species, *Culex nebulosus* is indicated by a grey line.

## Discussion

In this study, we analyzed the interplay between host community composition, habitat disturbance and virus prevalence in a multi-host and multi-taxa approach. We characterized the genetic diversity of RNA viruses in an entire family of hosts (*Culicidae*), which were sampled along an anthropogenic disturbance gradient. We subsequently studied prevalence patterns of all detected viruses per habitat type, as well as mosquito community composition per habitat type. We discovered an exceptionally high diversity of 49 distinct viruses, of which 34 were previously unknown members of seven different RNA virus families. We demonstrated that the majority of these viruses occurred at low minimum infection rates of 0.22 – 1.97 infected mosquitoes per 1000 tested mosquitoes. Nine viruses occurred more frequently across the disturbance gradient, of which five increased in prevalence from pristine to disturbed habitat types. We could show that the detection rates of these viruses corresponded to the abundance patterns of their specific mosquito host species. Importantly, the differences in virus prevalence were driven by the number of hosts present in a specific habitat type and not by changes in host infection rates. This interplay was named abundance effect. These data show that host community composition critically influences virus prevalence which has direct impact on our understanding of infectious disease emergence mechanisms.

The majority of the detected viruses grouped with insect-specific viruses (ISVs) in phylogenetic analyses, suggesting that vertebrates do not participate in the amplification and maintenance cycles of these viruses. However, the novel viruses Cimo peribunyavirus I and II formed a monophyletic clade that shared a most recent common ancestor with viruses of the genera *Orthobunyavirus* and *Pacuvirus*. Orthobunyaviruses are arboviruses that infect a great variety of vertebrates including humans (62). Pacuviruses were isolated from rodents and phlebotomine sandflies in Brazil (75, 76). Thus, the amplification cycle of Cimo peribunyavirus I and II may involve vertebrates and may be more complex than that of the other detected viruses. Further research is necessary to assess the host range of Cimo peribunyavirus I and II. Nevertheless, studying abundance patterns and geographic spread of insect-specific viruses in the light of habitat disturbance and shifts in host community composition can provide valuable insight into infectious disease dynamics. Mosquito-borne viruses are among the major global health concerns and the recent spread and epidemics caused by Zika and Chikungunya viruses have exemplified that there is an urgent need to understand viral emergence processes (26). Using mosquito-specific viruses as model viruses has many advantages as these viruses are more frequently found than arboviruses and no vertebrate host is required for virus maintenance and amplification which makes links between host community composition and virus abundance patterns easier to identify. Mosquitoes and their viruses are an ideal system to study such effects as this allows studying infectious disease dynamics in an entire family of hosts which can be tested in high numbers.

Our observed virus prevalence patterns are in agreement with several studies stating a heterogeneous effect of biodiversity on pathogen prevalence or disease risk (5, 22). Biodiversity can influence disease risk by different mechanisms like host regulation, changes in encounter rates or transmission rates, leading to an amplifying or diluting net effect (20). Areas with a high host richness likely harbor a high pathogen richness that might act as a source pool of novel diseases upon habitat change and increased contact to humans (2, 3). We observed the highest virus richness in intermediately disturbed and primary rainforest habitats supporting that biodiverse habitats are also rich in pathogens. However, highest viral prevalence rates were observed in villages and at camp sites, further supporting that intact or intermediately disturbed ecosystems also contain well-balanced spectra of viruses.

For those arboviruses, that are transmitted by a generalist mosquito species and use a competent host profiting from disturbance, a dilution effect can occur at local scales (7, 9). Several studies observed a dilution effect for West Nile virus (WNV) with either increasing non-passerine or total bird diversity (11, 77, 78) while others reported no protective effect of avian species richness on WNV prevalence (17, 79). For tick-borne encephalitis virus a dilution effect with increasing density of incompetent deer hosts was observed at local scale (80). Likewise, the prevalence of the directly transmitted hantavirus sin nombre virus is reduced at sites with higher rodent diversity as the persistence of the main host species (deer mouse) is reduced at diverse sites (12). A similar pattern was observed in our sampling for the five viruses GOLV, HEBV, FERV, Spilikins virus and CAVV belonging to four different families (*Phenuiviridae, Peribunyaviridae, Phasmaviridae* and *Mesoniviridae*). All these viruses used mosquitoes of the species *Culex nebulosus* as primary host which seemed to profit from disturbance and increased in abundance in disturbed versus pristine habitats (38% vs 3%). The prevalence of the five associated viruses was reduced in the diverse habitat types where the abundance of incompetent mosquito hosts increased and *Culex nebulosus* were rarely found. It is striking that this effect was observed for five taxonomically different viruses infecting the same host species. This effect was not observed for closer related viruses using other mosquito species as host which did not increase in abundance in disturbed habitats. These data suggest that the adaptability of mosquito hosts to changed environments plays a more important role for the increase in prevalence of associated viruses than the phenotypic or genetic characteristics of these viruses.

Contrary effects can be caused by scale-dependent effects or depending on additive or substitutive community assemblies (16). Additionally, the most competent host can either increase or decline with disturbance leading either to a dilution or amplification effect, respectively (7). In our investigated sample set, two viruses (Cimo phenuivirus II, family *Phenuiviridae*, and Cimo flavivirus I, family *Flaviviridae*) with higher prevalence in pristine than in disturbed habitats were detected. As observed for the five viruses mentioned above (GOLV, HEBV, FERV, Spilikins virus and CAVV), virus prevalence rates were closely linked to abundance rates of host mosquito species. Cimo phenuivirus II and Cimo flavivirus I were associated with *Uranotaenia* and *Coquillettidia* mosquitoes that declined in abundance in disturbed habitat types. An explanation why such an effect is rarely found could be that model pathogens selected in the field of disease ecology are usually zoonotic pathogens that spill over to humans in disturbed habitats and are known to cause outbreaks. Thus, there may exist a bias towards those pathogens that cause problems in disturbed habitat types. Pathogens that are rarely found in disturbed habitats may never be selected for disease ecology studies. These examples underline why an unbiased multi-host and multi-pathogen approach is crucial to uncover host community effects on pathogen prevalence patterns.

In other studies focusing on plant, vector-borne, indirectly and directly transmitted pathogens, the composition of the host community rather than total biodiversity was a predictor for pathogen distribution (5, 21). This is in agreement with the strong observed association between the prevalence of a certain virus and the abundance of its main mosquito host species in our study.

We detected novel strains of two previously characterized viruses, PCLV and Anopheles flavivirus, in *Aedes aegypti* and *Anopheles gambiae* mosquitoes, respectively, corresponding to previous findings of these viruses in these mosquito species in Asia and Africa (60, 66, 81). This further supports the specific associations between certain viruses and mosquitoes on a broader geographical scale.

Mosquitoes harbour a diverse natural virome that can influence subsequent virus infections (82, 83). We observed four frequently co-occurring viruses in *Culex nebulosus* mosquitoes (GOLV, HEBV, FERV and CAVV) that also grew together in cell culture, while in contrast the two *Culex decens*-associated viruses, Cimo rhabdovirus I and JONV, were rarely detected together. This could hint on the one hand at synergistic interactions and on the other hand at superinfection exclusion between different insect-specific viruses in mosquitoes. For several ISVs in the genera *Flavivirus* and *Alphavirus* an interference with related arboviruses was observed (84-86). A better knowledge of the virome of different mosquito species as well as studies on mutual interference might help to assess vector competence and possibilities of geographic spread.

Virus detection in mosquito homogenates independent of virus isolation extents potential virus findings but has the limitation that exogenous and integrated viruses can be discovered. The detection of likely integrated sequences derived from flavi- and rhabdoviruses in our mosquito sample is in agreement with frequent previous findings of NIRVS from these families in mosquito genomes (87-90). NIRVS are closer related to ISVs than to arboviruses probably limiting their influence on vector competence. The transovarial transmission of ISVs might increase the chance of germline integrations (72, 91). NIRVS can be transcriptionally active and might play a role in immunity against related viruses by producing piRNAs (87, 89, 90, 92). Collectively, our data show that only some viruses of a huge viral community benefited from ecosystem disturbance. This effect was found for five viruses (GOLV, HEBV, FERV, Spilikins virus and CAVV) and was determined by abundance patterns of their mosquito hosts which profited from habitat disturbance and strongly increased in numbers from pristine to disturbed habitats. In contrast, we also detected two viruses (Cimo phenuivirus II and Cimo flavivirus I) which were associated with mosquito hosts specific to primary habitat types. Detection rates of the associated viruses were also closely linked to host abundance and these viruses were found with higher frequencies in the primary and secondary rainforest. However, our analyses have also shown that virus prevalence is not always a consequence of the abundance of host species in a certain area. The prevalence pattern of JONV did not follow the abundance pattern of *Culex decens* mosquitoes and its abundance seems to be determined by unknown mechanisms, possibly by superinfection exclusion mediated by Cimo rhabdovirus I. The study of prevalence patterns of a broad genetic diversity of RNA viruses and their associated hosts allowed us to seek for general mechanisms influencing emergence and geographic spread of mosquito-associated viruses. No general dilution or amplification effect was observed. Instead, we identified changes in mosquito community composition causing an increase or decrease of the main host species as the most important factor determining virus prevalence patterns, an effect named abundance effect. Remarkably, host infection rates were not affected by higher host abundance in our study.

Taken together, our data show that host species composition is critical for virus abundance. Environmental changes that lead to an uneven host community composition and to more individuals of a single species is a key driver of virus emergence.

## Acknowledgements

We thank the Ivorian Ministry of Environment and Forest, the Ministry of Research and the directorship of the Taï National Park for the opportunity to conduct this research. We thank all field assistants for help in the field and the Taï chimpanzee project and Fabian Leendertz for logistic support during field work. We are grateful to Verena Heyde, Christian Hieke and Friederike Schröder for excellent laboratory assistance. The work was funded by the Federal Ministry of Education and Research (BMBF) under project number 01KI1716 as part of the Research Network Zoonotic Infectious Diseases and by the Deutsche Forschungsgemeinschaft under project B02 as part of the collaborative research center CRC228.

## Author contribution statement

Conceptualization, S.J. and K.H.; Investigation, K.H., M.M., F.Z. and A.K.; Formal Analysis, S.K.S and K.H.; Resources, S.J.; Supervision, S.J.; Funding Acquisition, S.J.; Writing - Original Draft Preparation, K.H. and S.J.; Writing - Review & Editing, all authors.

## Supplementary Information

**SI Table 1:** Model estimates (base: habitat Primary forest and mosquito group *Anopheles* sp.) for the counts of mosquito individuals per mosquito group and habitat. Significant predictors show the difference to the baseline combination. CulAnn: *Culex annuloris*, CulDec: *Culex decens*, CulNeb: *Culex nebulosus*, Cul_spec: other *Culex* species; Ura_spec: *Uranotaenia* species; Coq_spec: *Coquillettidia* species; Ano_spec: *Anopheles* species; others: all other grouped species. PF: primary forest, SF: secondary forest, A: agriculture, C: camp, V: village.

**SI Table 2:** Significant differences in the infection probability with the most abundant viruses in the different habitats (only Tukey contrasts are shown; bold: p < 0.05; italic: p < 0.1). No significant differences were detected for FERV (n = 20), JONV (n = 17), Spilikins virus (n = 12) and Cimo flavivirus I (n = 13; no findings in camp and village). Combinations with zero virus detections shown in light grey. All virus detections were positively associated with the number of mosquitoes per pool (model estimates not shown).

**SI Figure 1:**
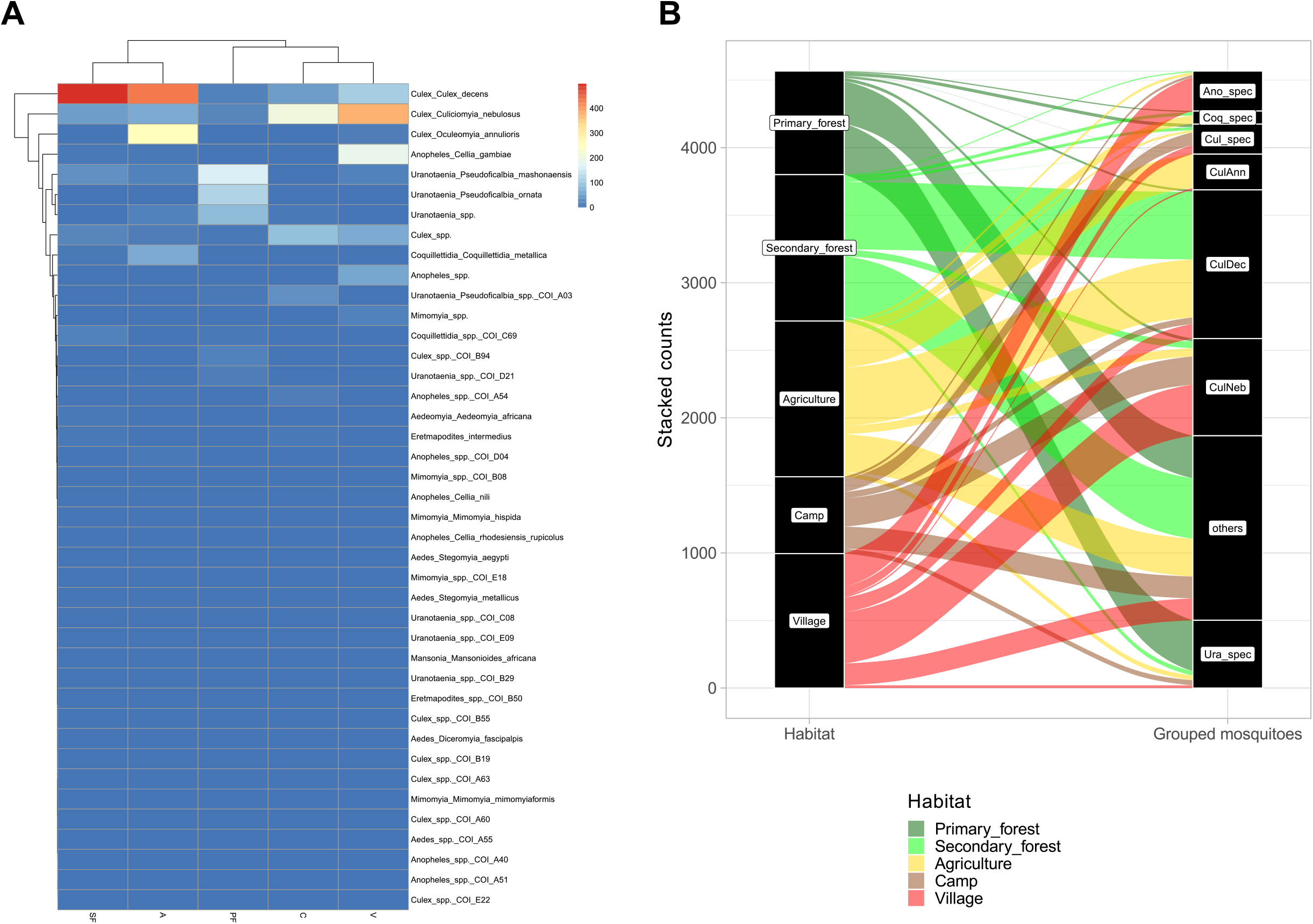
Biodiversity analyses. (A) Hierarchical cluster analysis based on associations between mosquito species. Main mosquito groups were identified as CulAnn, CulDec, CulNeb, Cul_spec, Ura_spec, Ano_spec and Coq_spec. (B) Alluvial plot showing the distribution of the main mosquito species groups to the respective habitats. CulAnn: *Culex annulioris*, CulDec: *Culex decens*, CulNeb: *Culex nebulosus*, Cul_spec: other *Culex* species; Ano_spec: *Anopheles* species; Ura_spec: *Uranotaenia* species; Coq_spec: *Coquillettidia* species, others: all other grouped species. PF: primary forest, SF: secondary forest, A: agriculture, C: camp, V: village.

**SI Figure 2:**
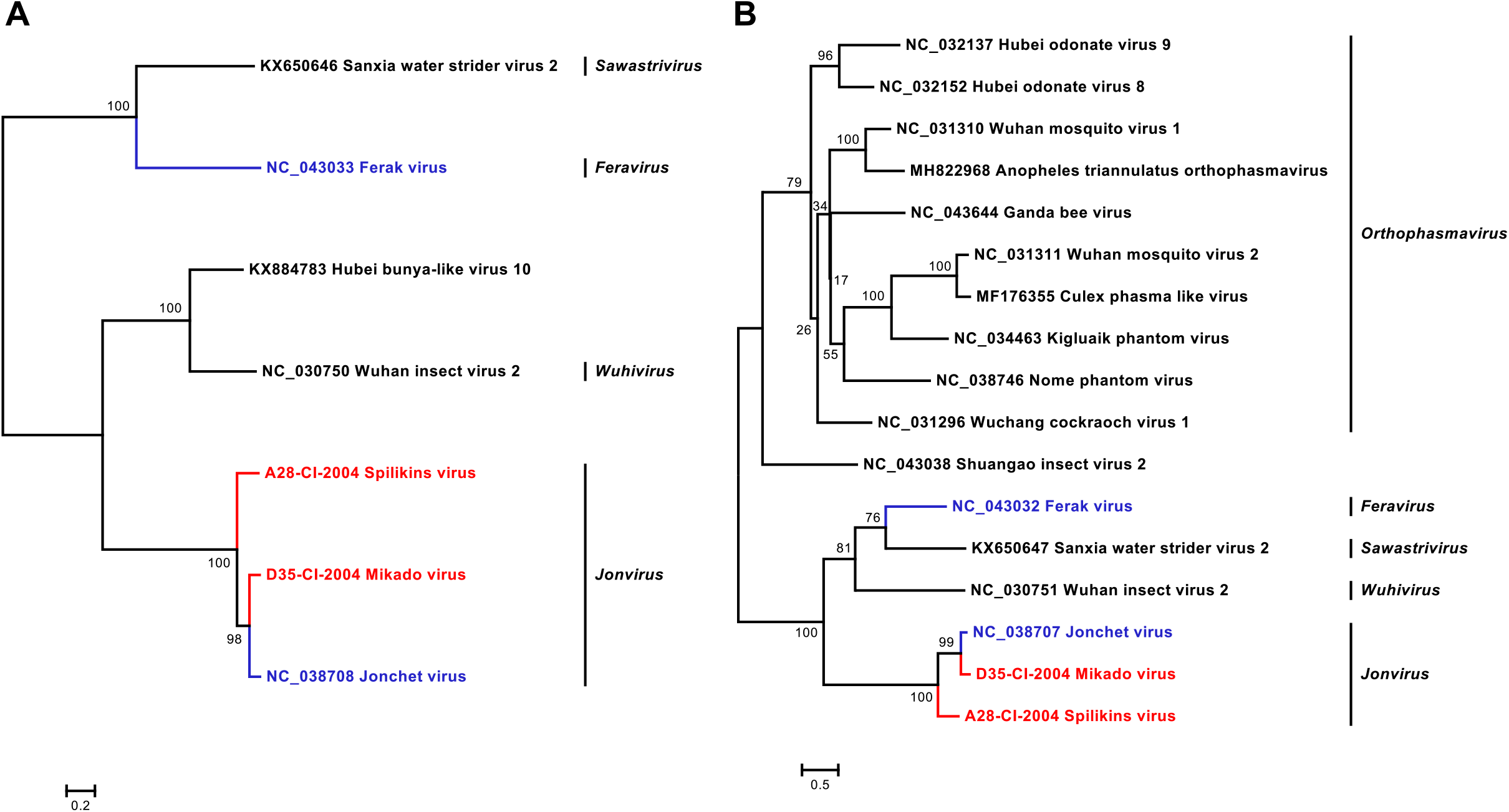
Phylogenetic analyses of detected phasmaviruses. Phylogenetic trees were inferred with PhyML (WAG (A) and LG (B) substitution model) based on MAFFT-E protein alignments covering the glycoproteins (**A**) and nucleocapsid (**B**) proteins of the family *Phasmaviridae*. Novel viruses from this study are indicated in red and previously published viruses detected in our data set are indicated in blue.

**SI Figure 3:**
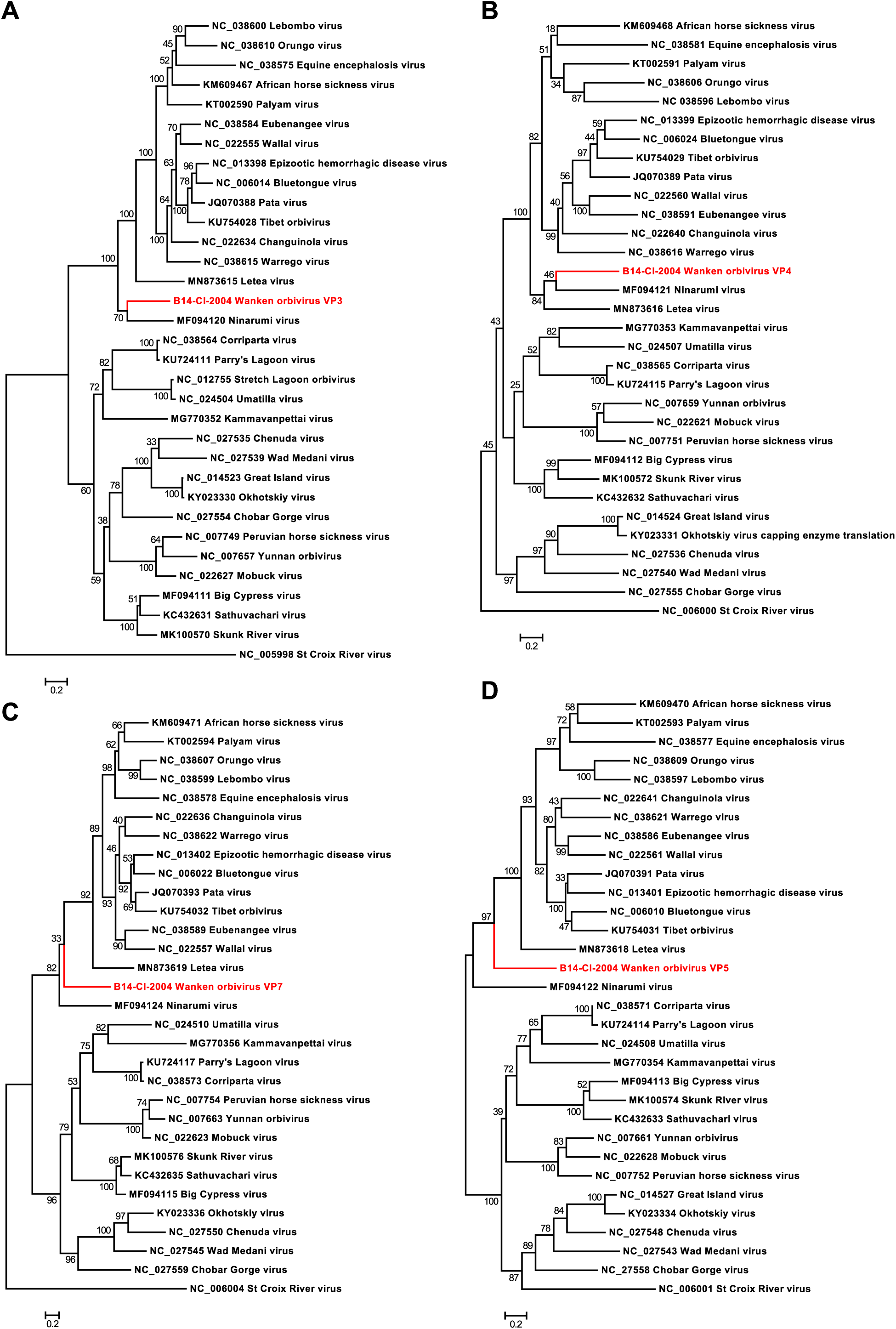
Phylogenetic analyses of the detected WKOV and members of the genus *Orbivirus*. Phylogenetic trees were inferred with PhyML (LG substitution model) based on MAFFT-E protein alignments of VP3 (**A**), VP4 (**B**), VP7 (**C**) and VP5 (**D**). WKOV is indicated in red.

**SI Figure 4:**
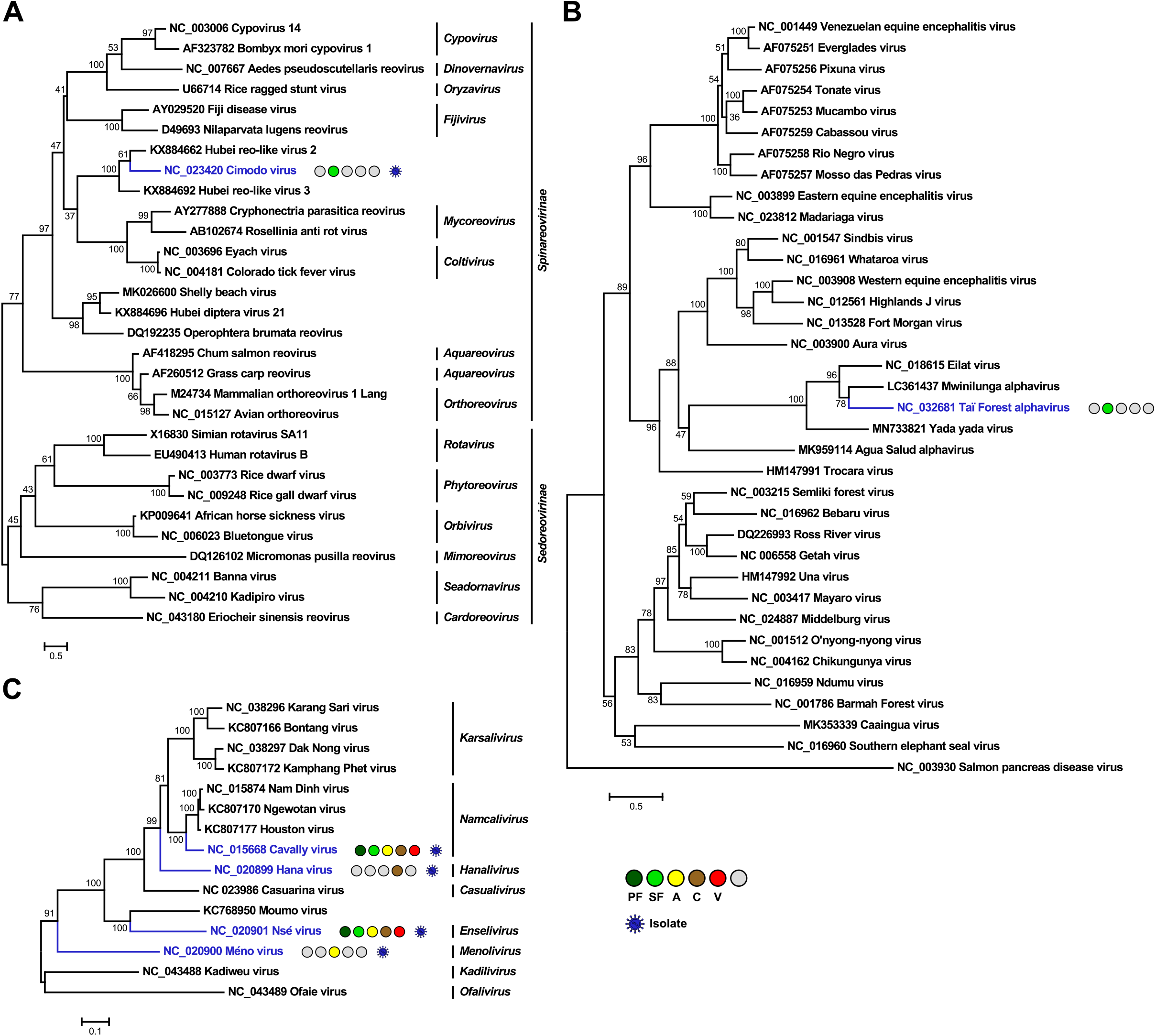
Phylogenetic analyses of detected reoviruses, alphaviruses and mesoniviruses. Phylogenetic trees were inferred with PhyML (LG substitution model) based on a MAFFT-E protein alignment of the polymerase of members of the family *Reoviridae* (**A**) or with PhyML (GTR substitution model) based on MAFFT-E nucleotide alignments covering the E2-6K-E1 region of members of the genus *Alphavirus* (**B**) of the polymerase of members of the family *Mesoniviridae* (**C**). Novel viruses from this study are indicated in red, previously published viruses detected in our data set are indicated in blue and detected virus-like sequences are indicated in grey. Sample origin from the different habitat types is indicated by colored circles while no detection is indicated by grey circles. Live virus isolates are marked with a blue virion. Abbreviations are PF, primary forest; SF, secondary forest; A, agriculture; C, camp and V, village.

**SI Figure 5:**
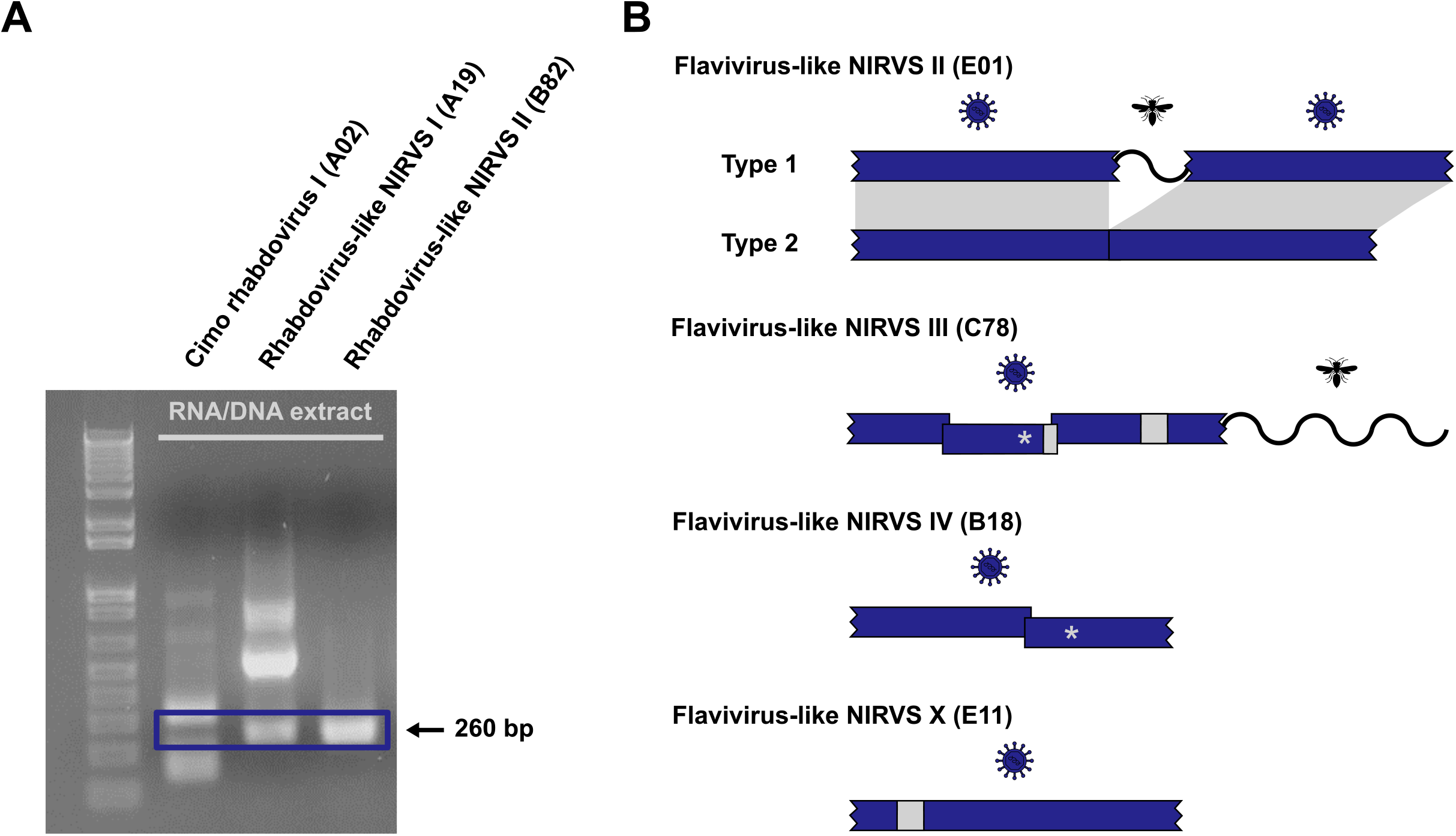
Potential NIRVS. (**A**) PCR amplicons of the generic rhabdovirus PCR assay. RNA/DNA extracts without reverse transcription were used for the PCR. Nested PCR amplicons of Cimo rhabdovirus I and two rhabdovirus-like NIRVS were visualized by ethidium bromide-stained agarose gel electrophoresis. Amplicons with the expected size of 260 bp are framed by a blue box. (**B**) Schematic representation of selected flavivirus-like NIRVS. Stop codons are indicated by an asterisk, deletions are shown as light grey boxes, frame shifts are indicated by overlapping blue boxes and insertions with similarity to insect genes are shown as waved lines.

**SI Figure 6:**
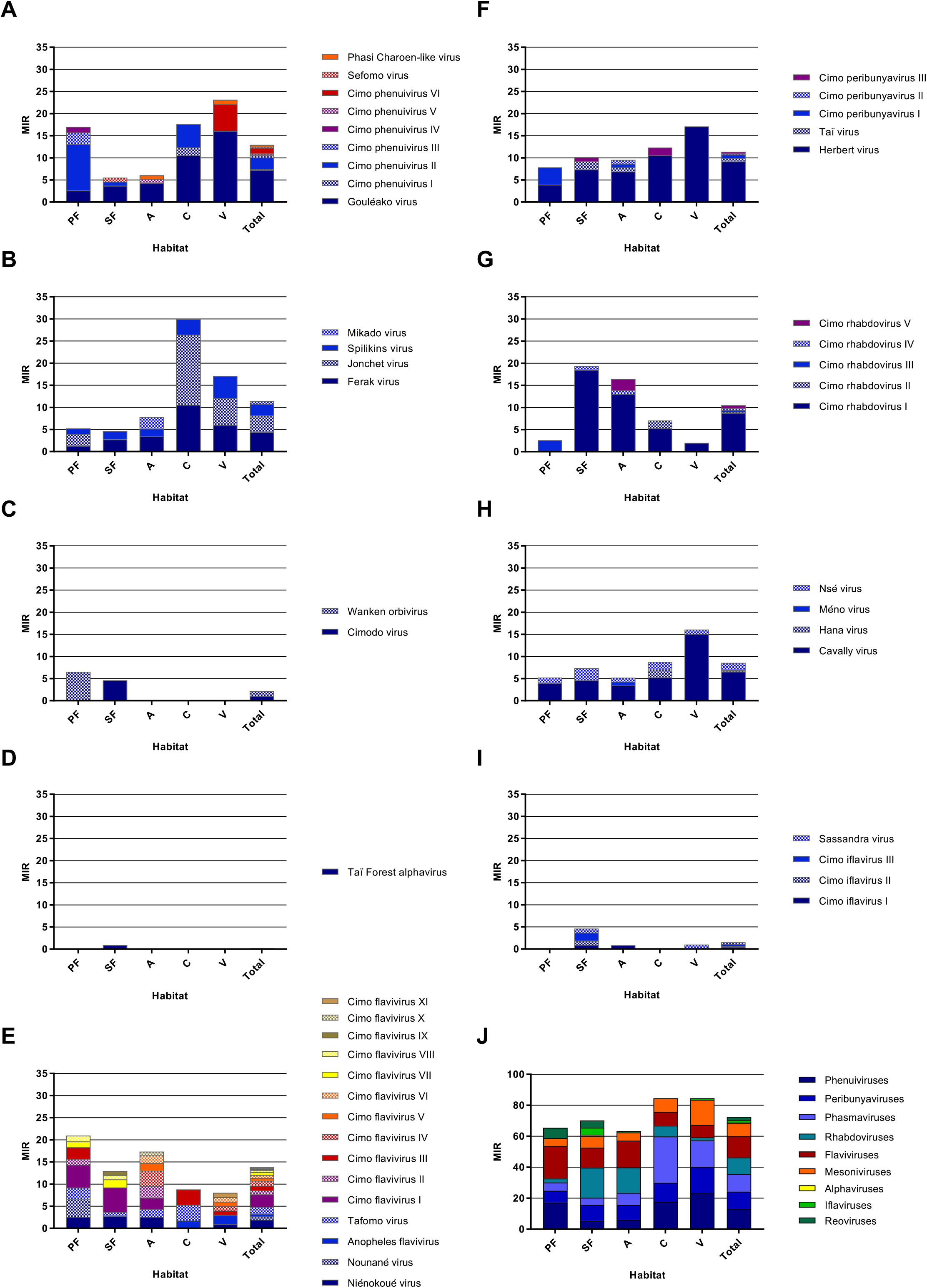
Cumulative MIR per virus taxon. The cumulated MIR per 1000 mosquitoes was calculated for the analyzed taxa *Phenuiviridae* (**A**), *Phasmaviridae* (**B**), *Reoviridae* (**C**), *Alpha-virus* (**D**), *Flavivirus* (**E**), *Peribunyaviridae* (**F**), *Rhabdoviridae* (**G**), *Mesoniviridae* (**H**), and *Iflaviridae* (**I**), as well as for all detected viruses (**J**) in the different habitat types and for the complete data set. The different viruses or taxa are shown in different colours.

